# Genome-wide analysis reveals a switch in the translational program upon oocyte meiotic resumption

**DOI:** 10.1101/685594

**Authors:** Xuan G. Luong, Enrico Maria Daldello, Gabriel Rajkovic, Cai-Rong Yang, Marco Conti

**Affiliations:** Center for Reproductive Sciences, University of California, San Francisco, CA 94143, USA; Eli and Edythe Broad Center of Regeneration Medicine and Stem Cell Research, University of California, San Francisco, CA 94143, USA; Department of Obstetrics and Gynecology and Reproductive Sciences, University of California, San Francisco, CA 94143, USA.

**Keywords:** Meiosis, RNA-Seq, mouse oocyte, translation, CPEB1

## Abstract

During oocyte maturation, changes in gene expression depend exclusively on translation and degradation of maternal mRNAs rather than transcription. Execution of this translation program is essential for assembling the molecular machinery required for meiotic progression, fertilization, and embryo development. With the present study, we used a RiboTag/RNA-Seq approach to explore the timing of maternal mRNA translation in quiescent oocytes as well as in oocytes progressing through the first meiotic division. This genome-wide analysis reveals a global switch in maternal mRNA translation coinciding with oocyte re-entry into the meiotic cell cycle. Messenger RNAs whose translation is highly active in quiescent oocytes invariably become repressed during meiotic re-entry, whereas transcripts repressed in quiescent oocytes become activated. Experimentally, we have defined the exact timing of the switch, the repressive function of CPE elements, and identified a novel role for CPEB1 in maintaining constitutive translation of a large group of maternal mRNAs during maturation.

## Introduction

Cell development relies on elaborate changes in gene expression in order to transition through different phenotypic and functional stages that ultimately lead to terminal differentiation. Changes in gene expression are achieved through transcriptional and post-transcriptional regulations. Although transcriptional regulation is understood in considerable detail(Chen and Dent, 2014; Klemm et al., 2019), much less is known about the molecular machinery involved in translation regulation.

Large oligomeric complexes involving proteins and non-coding RNAs are assembled on the mRNA (Rissland, 2017) to regulate its interaction with ribosomes, its translation rate, and its stability (Rissland et al., 2017; Wu and Brewer, 2012). In somatic cells, numerous observations indicate that translation is intimately coupled with degradation of mRNAs (Jonas and Izaurralde, 2015; Wu and Brewer, 2012). Proteins recruited to the mRNA interact with elements located throughout the length of the transcript (Cheng et al., 2017; Rissland, 2017). However, complexes nucleated around the 5’ and 3’ untranslated regions (UTRs) play a predominant role in translation and stabilization, often by controlling the length of the poly(A) tail, which is present in most mRNAs (Jacobson and Peltz, 1996; Rissland et al., 2017). In gametes and embryos, particularly, the poly(A) tail determines the translation rate and stability of a mRNA (Clarke, 2012; Radford et al., 2008; Richter and Lasko, 2011; Subtelny et al., 2014; Tay et al., 2000; Yartseva and Giraldez, 2015).

Germ cells are unique in their properties as they progressively acquire specialized function during development (Clarke, 2012). At the same time, they maintain traits that allow for rapid transition to totipotency (Seydoux and Braun, 2006). Throughout development, germ cells often rely on unique post-transcriptional regulations rather than on transcription itself (Clarke, 2012; Kimble and Crittenden, 2007). Striking examples of this property are the growth and maturation stages of an oocyte and its transition to zygote and early embryo (Clarke, 2012; Yartseva and Giraldez, 2015). During the growth phase, oocytes amass large number of maternal mRNAs through high transcriptional activity. These mRNAs are either used immediately to synthesis proteins involved in growth or are stored for future use. Indeed in all species studied, transcription ceases when an oocyte is fully grown and resumes only in the embryo. Thus, critical steps in oocyte maturation and early embryo development rely exclusively on a program of maternal mRNA translation.

Some properties of the molecular machinery involved in maternal mRNA translation repression or activation have been elucidated in model organisms (Kimble and Crittenden, 2007; Tadros and Lipshitz, 2005; Yartseva and Giraldez, 2015). In frogs, the cytoplasmic polyadenylation element-binding protein (CPEB) is considered a master regulator of polyadenylation and translation (Mendez and Richter, 2001; Richter, 2007). Much less is known about the role of CPEB in mammalian oocytes. Here, we have used a genome-wide approach to investigate the role of this RNA-binding protein (RBP) during the transition from quiescence to re-entry into the meiosis. Through a detailed time course, we have investigated the temporal association between maternal mRNA translation and the different steps involved in oocyte re-entry into and progression through the meiosis. Using a RiboTag/RNA-Seq strategy, we describe a genome-wide switch in the translation program of maternal mRNAs, and define new, critical functions of CPEB in the control of this switch.

## Results

### Re-entry into meiosis coincides with rapid translational changes of stable mRNAs

We have used a RiboTag/RNA-Seq strategy to characterize mRNA translation in oocytes arrested at prophase I (GV) and in those undergoing meiotic maturation. (Fig. 1a). This strategy has been previously validated (Martins et al., 2016; Yang et al., 2017) and additional quality controls are reported here (Supplementary Fig. 1a-d). Upon meiotic resumption, there were both progressive increases and decreases in ribosome loading of maternal mRNAs (Fig. 1b). By late metaphase I (MI), mRNAs were either constitutively translated (n = 4284, CONSTITUTIVE), translationally repressed (n = 1722, DOWN), or translationally activated (n = 1537, UP) (FDR ≤ 0.05, Supplementary Fig. 1e). Total mRNA levels remained stable up to MI and significant destabilization was detectable only for 3% of maternal mRNAs at late MI (Fig. 1b and Supplementary Fig. 1f). Comparison of changes in total mRNA levels (transcriptome) to changes in ribosome-associated mRNA levels (translatome) confirms this late MI destabilization (Fig. 1c), which became prominent later during MII arrest, with a subset of mRNAs remaining stable even if their translation was repressed. Changes in translation that initiate during MI were extended into and amplified at MII (Fig. 1d). Furthermore, there is a strong, positive correlation between translational changes at late MI and MII (Supplementary Fig. 1g). Therefore, the patterns of differential ribosome loading are consistent across multiple *in vitro* and *in vivo* biological replicates and across two distinct detection platforms.

**Fig. 1.**
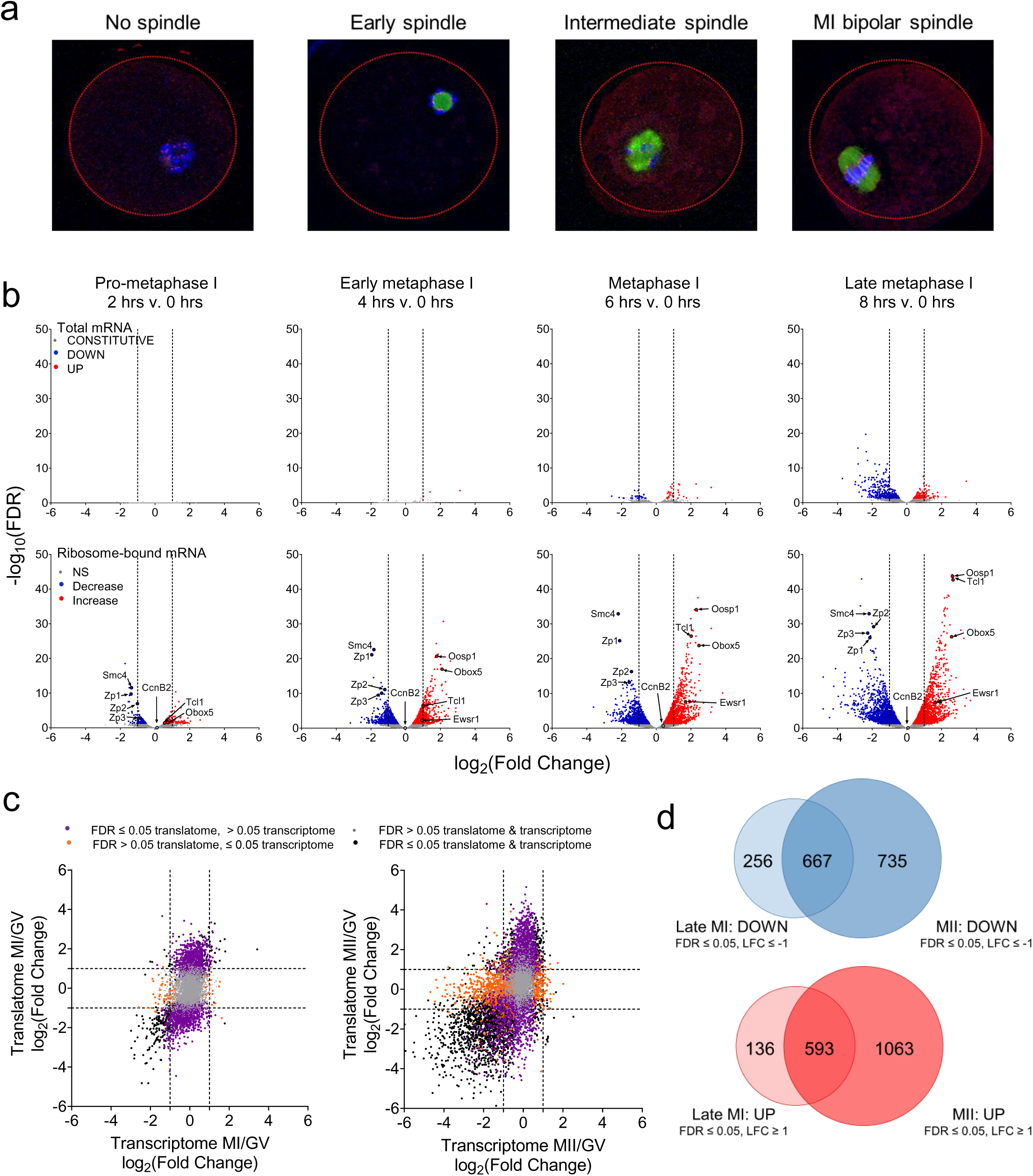
The translational program during oocyte meiotic cell cycle involves both translational repression and activation. **a)** Spindle and chromatin conformation in the oocyte during meiosis. Oocytes were matured *in vitro* and fixed at 0, 2, 4, 6, and 8 hrs post-meiotic resumption. Immunofluorescence staining for tubulin (green), kinetochores (red), and chromatin (blue) was performed. Maturing oocytes either presented chromosome condensation, but no spindle assembly (2 hrs, pro-metaphase I), visible initial spindle formation (4 hrs, early MI), progressive spindle formation with kinetochore attachment (6 hrs, MI), or a fully attached, MI-bipolar spindle (8 hrs, late MI). **b)** Total mRNA levels and differential ribosome loading during meiotic progression as compared to prophase I arrest. Oocytes were matured *in vitro* and collected at 0, 2, 4, 6, and 8 hrs post-meiotic resumption. Total RNA samples were collected prior to RiboTag-IP for each time point. cDNA libraries were prepared from total and ribosome-bound RNA samples, RNA-Seq was performed, and the data processed and analyzed as described in the “Methods”. The data are presented as volcano plots with log_2_(fold change) (LFC) CPM at each time point compared to 0 hrs and plotted against false discovery rate (FDR). Statistically significant increased (red) and decreased (blue) (FDR ≤ 0.05) genes are reported as well as non-significant changes (grey). LFCs ≤ −1 or ≥ 1 are considered biologically significant and are marked by dashed lines. Two biological replicates of 200 oocytes per time point were used for this experiment. **c)** Changes in the transcriptome and translatome during meiosis. The late MI data are derived from the experiment described in Fig. 1b, MII translation data are from a deposited dataset generated from oocytes matured *in vivo* followed by polysome fractionation/microarray (polysome array)(Chen et al., 2011; Conti and Franciosi, 2018); and prophase-to-MII total mRNA data were from a deposited dataset(Su et al., 2007). The data are reported as scatterplots with LFC in total mRNA CPM at either late MI or MII compared to prophase I arrest versus the LFC of ribosome-bound mRNA CPM at the same time points. We identified four groups of messages: transcripts that showed significant changes only in translation (purple), significant changes only in total message levels (orange), no significant changes in translation nor in total transcript levels (grey), and significant changes in both translation and total transcript levels (black); significant changes are defined as FDR ≤ 0.05. Two biological replicates of 200 oocytes per time point were used to generate the RNA-Seq data, while six biological replicates of 500 oocytes per time point were used to generate the polysome array data. **d)** Overlap of translatome changes between late MI and MII. Both DOWN (blue) and UP (red) genes were analyzed. The data were collected as described in Fig. 1c. LFCs ≤ −1 or ≥ 1 with FDR ≤ 0.05 are considered statistically significant.

We compared the maternal mRNAs that are translational repressed and those that are degraded in our dataset with those stabilized in mouse oocytes depleted of YTHDF2 (Ivanova et al., 2017), a RNA m(6)A reader, or CNOT6L (Horvat et al., 2018), a component of the CCR4 complex, and found little overlap (Supplementary Fig. 2a-d). We do, however, observe overlap between our data and those mRNAs that are stabilized in MII in BTG4^-/-^ mouse oocytes (Supplementary Fig. 2e and f), confirming that repressed mRNAs are eventually destabilized. BTG4 is a member of the TOB family of proteins that interacts with CNOT7/8 and is required for mRNA destabilization in MII (Liu et al., 2016; Pasternak et al., 2016; Yu et al., 2016).

### Divergent mechanisms control gene expression during mitosis and meiosis

Repression of maternal mRNA translation is associated predominantly with mitochondrial and ribosomal biogenesis, while translationally activated mRNAs code for proteins with functions related to cell cycle and embryo development (Conti and Franciosi, 2018). Gene ontology (GO) analysis of UP and DOWN transcripts at late MI reinforces this association (Fig. 2a). Genome-wide comparison of our RNA-Seq data to those available for translation and transcription during mitosis (Park et al., 2016) did not reveal any significant correlation (Fig. 2b), suggesting profound differences in gene expression regulation during these processes. When the comparison between mitosis and meiosis is restricted to genes specific to cell cycle function, only nine mRNAs overlap in translation activation between mitosis and meiosis (Fig. 2c). However, a sizable group of mRNAs whose translation is activated during meiosis instead is activated transcriptionally at the S-to-M-phase transition. Although limited, overlap is also detected when translation repression during meiosis is compared with changes in translation during mitosis. Manual curation of the data confirms that decreased translation of *Cdk1* and increased translation of *Bub1b* occur during both mitosis and meiosis (Supplementary Fig. 3).

**Fig. 2.**
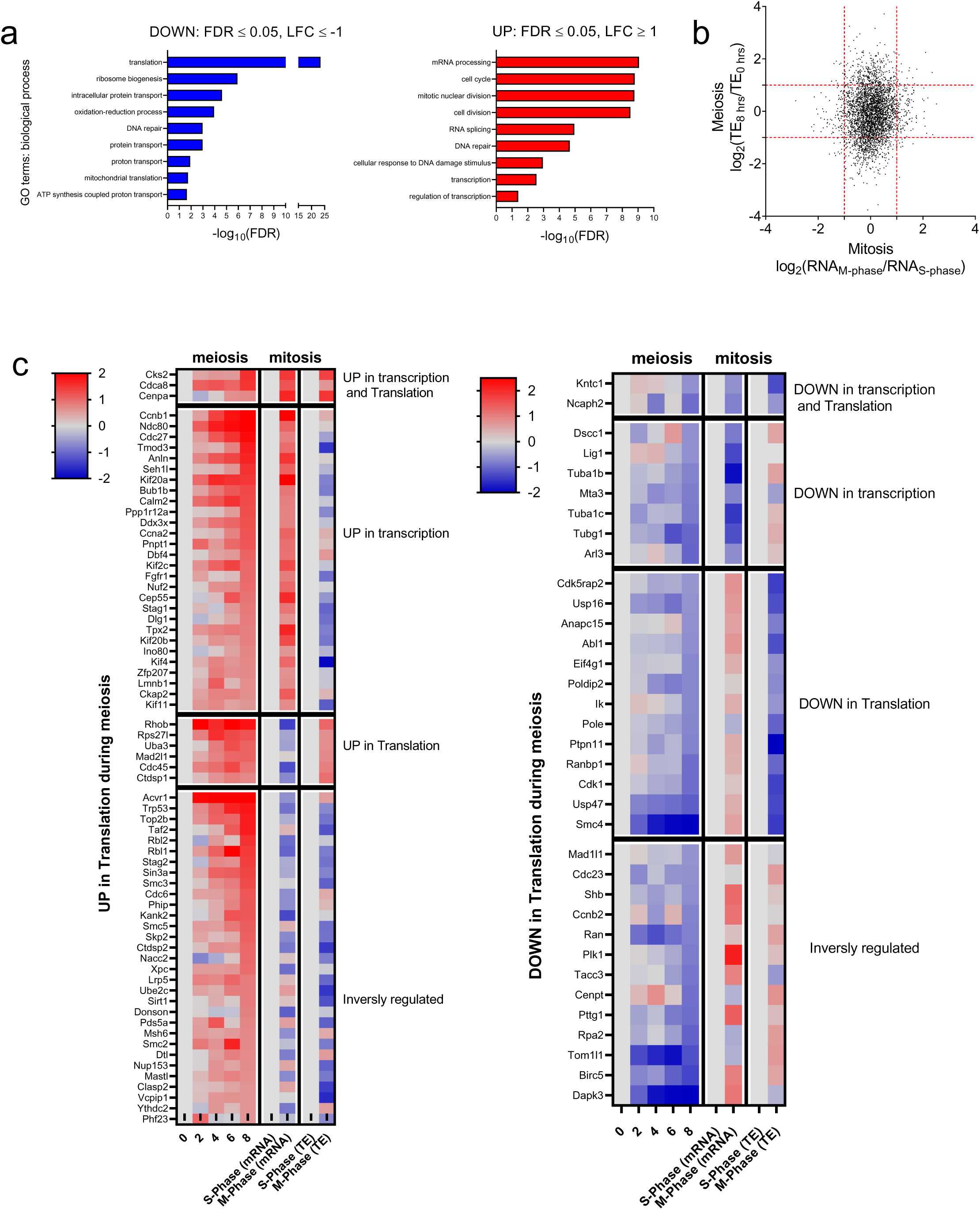
Cell cycle components are regulated via translation in meiosis, but via transcription in mitosis. **a)** Gene ontology analysis of DOWN and UP genes. DOWN (blue) and UP (red) mRNAs significantly changed from 0 hrs (prophase I arrest) to 8 hrs (late MI) post-meiotic resumption were used (−1 ≥ LFC ≥ 1 and FDR ≤ 0.05). Only terms with FDR ≤ 0.05 were considered. **b)** Pairwise comparison of translation during meiosis and transcription during mitosis. FCs in translational efficiency (TE) from 0 hrs (prophase I arrest) to 8 hrs (late MI) in our RNA-Seq dataset are plotted against FCs in RNA levels from S-phase to M-phase found in a deposited dataset(Park et al., 2016). **c)** Heat maps comparing fold changes in translation of cell cycle components during meiosis in oocytes and fold changes in transcription or translation during the mitotic cell cycle. The data were collected as described in Fig. 2b. Genes involved in the cell cycle are as defined under GO:0007049.

### The pattern of mRNA translation in GV-arrested oocytes predicts changes occurring during the G2-to-M-phase transition

During prophase I, maternal messages display a broad spectrum of translation efficiencies (TEs) (Fig. 3a), calculated as the ratio between ribosome-bound and total mRNA levels; there is a seven-fold difference when comparing the average TE of the 10% of mRNAs with the highest TEs (high-TEs) to that of the 10% with the lowest TEs (low-TEs). To validate that TE reflects rate of translation, we related these values to other available measurements (Morgan et al., 2017; Wang et al., 2010) in GV-arrested oocytes. High-TE messages have significantly longer poly(A) tails (≥ 70 nts) (Fig. 3b) and are associated with increased protein accumulation as assessed by mass spectrometry (Fig. 3c).

**Fig. 3.**
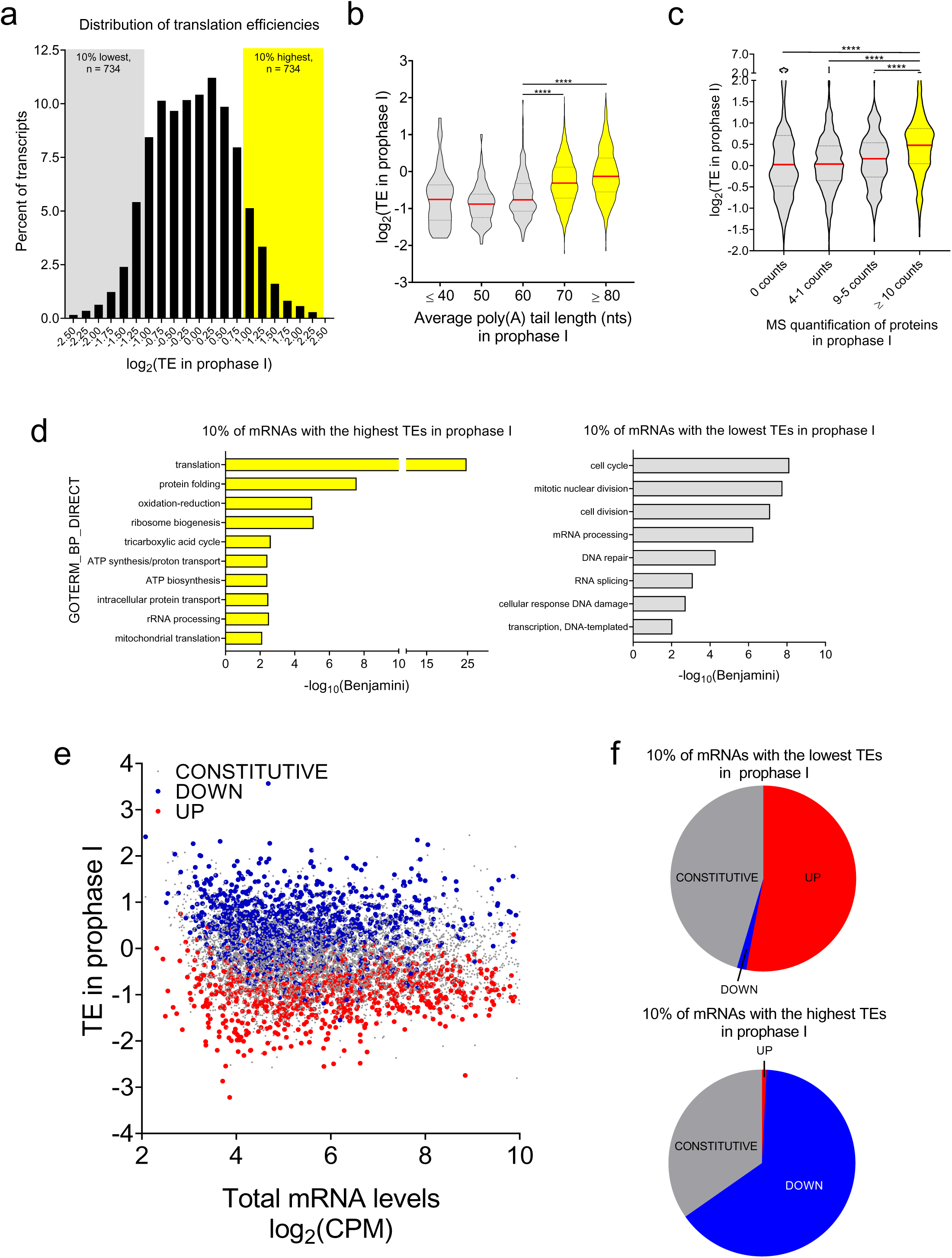
Genome-wide analysis of translation reveals a switch in the translation program at the quiescence-to-meiotic cell cycle re-entry transition. **a)** Histogram of translational efficiencies in GV-arrested oocytes. Translation efficiency (TE) for individual mRNAs was calculated as the ratio between ribosome-associated and total mRNA CPMs. Plotted is the distribution of TE values of maternal mRNAs during prophase I arrest. The 10% of mRNAs with the lowest TEs are designated as low-TE mRNAs (n = 734, grey box) and the 10% of mRNAs with the highest TEs as high-TE mRNAs (n = 734, yellow box); this definition is used for all the subsequent comparisons. **b)** Genome-wide relationship between TE and poly(A) tail length in GV-arrested oocytes. TE was calculated for individual mRNAs as described in Fig. 3a. Deposited TAIL-Seq data on poly(A) tail length of maternal mRNAs during prophase I arrest(Morgan et al., 2017) were associated with their TEs during this time. Median values are represented by red lines and the 25 and 75% quartiles are represented by black, dashed lines. Statistical significance was evaluated by unpaired, two-tailed t-tests. ****: p < 0.0001. **c)** Genome-wide relationship between TE and protein levels in GV-arrested oocytes. TE was calculated for individual mRNAs as described in Fig. 3a. Deposited data on protein levels in GV-arrested oocytes as quantified by mass spectrometry(Wang et al., 2010) were associated with the TEs of maternal mRNAs. Median values are represented by red lines and the 25 and 75% quartiles are represented by black, dashed lines. Statistical significance was evaluated by unpaired, two-tailed t-test; ****: p < 0.0001. **d)** Gene ontology analysis of low- and high-TE maternal mRNAs in GV-arrested oocytes. Only terms with a Benjamini coefficient ≤ 0.05 were considered. **e)** Genome-wide relationship between TE in prophase I arrest and translation pattern during meiotic resumption of oocyte maternal mRNAs. TE was calculated for individual mRNAs as described in Fig. 3a. The data are presented as a scatterplot of total mRNA CPMs compared to TE values in GV-arrested oocytes. Transcripts are then categorized as CONSTITUTIVE (grey), DOWN (blue), or UP (red) on the basis of their translation pattern during maturation to late MI (−1 ≥ LFC ≥ 1 and FDR ≤ 0.05). **f)** Detailed analysis of the relationship between TE in prophase I arrest and translation pattern during meiotic resumption of low- and high-TE mRNAs. Pie charts report the percentage of low- or high-TE mRNAs in GV-arrested oocytes that are UP, DOWN, or CONSTITUTIVE during meiotic maturation. Ninety-nine percent of low-TE mRNAs are either UP (53%) or CONSTITUTIVE (46%) and all the high-TE mRNAs are either DOWN (65%) or CONSTITUTIVE (35%).

GO analysis of low- and high-TE messages during prophase I arrest revealed associations antithetical to those found during meiotic maturation (Fig. 2a). Functions important for oocyte growth are significantly enriched for high-TE mRNAs, while functions important during oocyte maturation are enriched in low-TE mRNAs (Fig. 3d). We hypothesize that, upon meiotic resumption, a switch in the translation program occurs in order for the oocyte to progress through meiosis and prepare for embryogenesis. Indeed, transcripts with greater TEs become translationally downregulated (DOWN), while transcripts with lower TEs become activated (UP) during meiotic maturation (Fig. 3e). More detailed analysis reveals that translation of 99% of low- TE mRNAs is constitutive or upregulated during meiotic maturation, while translation of virtually all of high-TE mRNAs is constitutive or repressed (Fig. 3f).

### Unique mRNA features are associated with the opposing translation patterns in GV-arrested oocytes

To understand how such a broad array of TEs is established in GV-arrested oocytes, we correlated these values with various mRNA features genome-wide (Fig. 4a). ATG density, GC content, and length of the 5’UTR are inversely correlated with TE. As for the 3’UTR, polyadenylation signal (PAS) density and GC content are positively correlated, whereas DAZL-binding element density, 3’ UTR length, and cytoplasmic polyadenylation element (CPE) density are all inversely correlated. Detailed analysis confirms this strong inverse relationship between CPE density and TE, as 87% of low-TE mRNA contain putative CPEs in the 3’UTR, while this is only true for 57% of high-TE mRNAs (Fig. 4b). The absence of CPEs proximal to the PAS is significantly associated with greater TEs (Fig. 4c), and the closer a CPE is to the PAS, the less efficiently a message is translated during prophase I arrest (Fig. 4d).

**Fig. 4.**
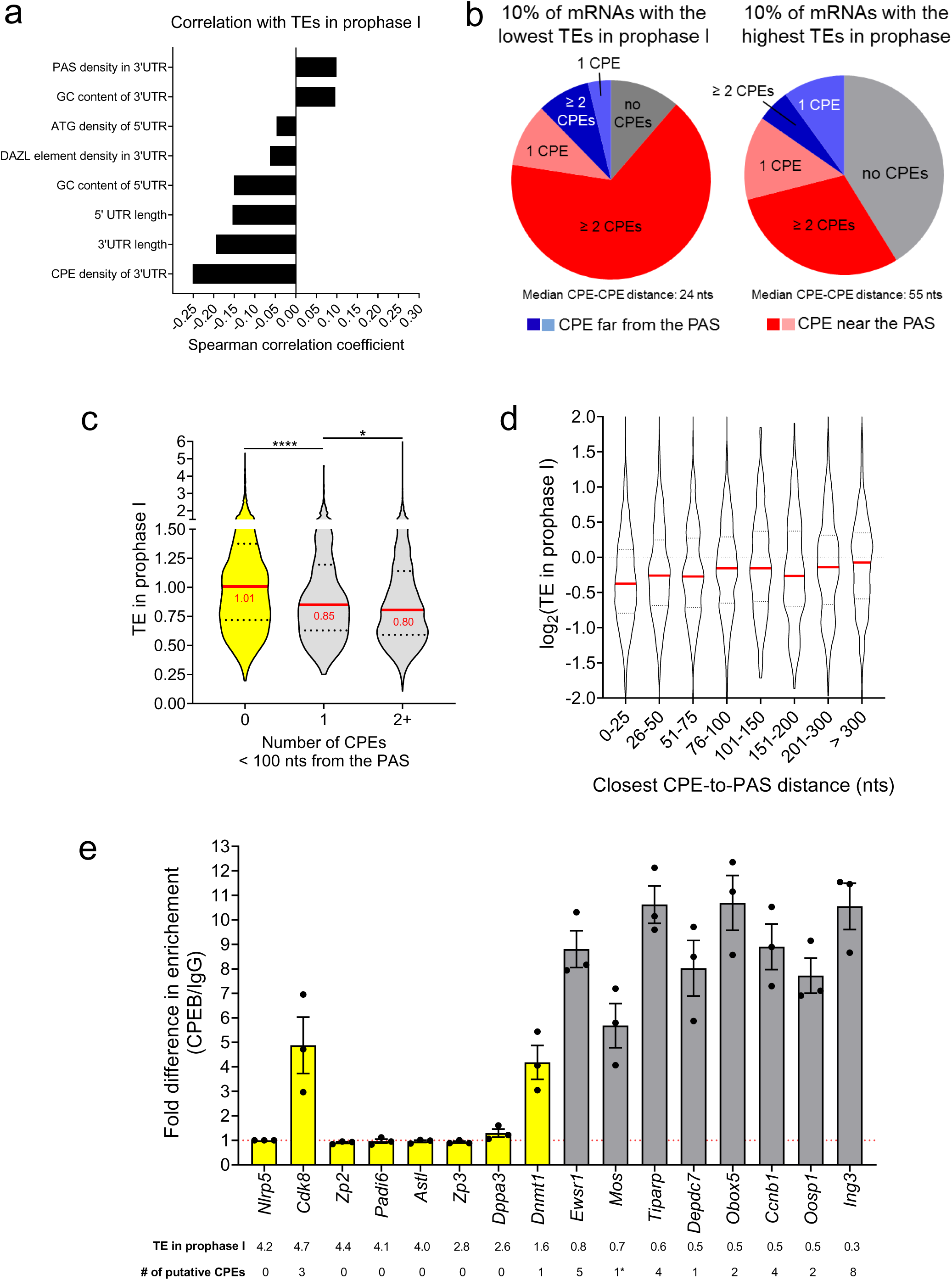
Features associated with maternal mRNAs translated with high or low efficiency in GV-arrested oocytes. **a)** Genome-wide correlation between mRNA features with TE in GV-arrested oocytes. PAS density in the 3’UTR, GC content in the 3’ and 5’UTRs, ATG density in the 5’UTR, DAZL and CPEB *cis*-acting element densities in the 3’UTR, and 3’UTR and 5’UTR lengths were calculated as detailed in the “Methods”. These data were then correlated with TEs during prophase I arrest and Spearman correlation coefficients were calculated for every comparison; p < 0.0001 for all pairs. In mRNAs with higher TEs, the reduced number of 3’UTR *cis*-acting elements is not due to shorter 3’UTR length, as element number was normalized for 3’UTR length when calculating densities. **b)** Detailed analysis of the relationship between TE in prophase I arrest and presence of CPEs in the 3’UTR of low- and high-TE mRNAs. Pie charts report the percentage of low- or high-TE mRNAs in GV-arrested oocytes where a CPE could be identified. Scanning for CPE in the 3’ UTRs was performed as detailed in the “Methods”. **c)** Genome-wide relationship between TE and number of CPEs found within 100 nts of the PAS in GV-arrested oocytes. Median values are represented by red lines and the 25 and 75% quartiles are represented by black, dashed lines. Statistical significance was evaluated by unpaired, two-tailed t-tests; *: p = 0.0313; **** p < 0.0001. **d)** Detailed analysis of the relationship between TE in prophase I arrest and the distance of the closest CPE to the PAS. Median values are represented by red lines and the 25 and 75% quartiles are represented by black, dashed lines. **e)** Enrichment of low-TE mRNAs bound to CPEB1 in GV-arrested oocytes. GV-arrested oocytes were collected and RNA-IP followed by RT-qPCR was performed as described in the “Methods”. *Nlrp5* was used as a reference gene as it is known to not bind to CPEB1. Three biological replicates of 200 oocytes per time point were used and RT-qPCR reactions were run in triplicate. Data are presented as fold difference in mRNA levels in CPEB1-IP as compared to the IgG-IP. The bars represent the mean ± SEM of three experiments. TE and the number of putative CPEs for each gene are reported. *The *Mos* 3’UTR has a single embryonic CPE(Simon and Richter, 1994).

This correlation was confirmed by CPEB1 RNA-immunoprecipitation (RNA-IP) followed by RT-qPCR (Fig. 4e). While only two of the eight candidate high-TE mRNAs (*Cdk8* and *Dnmt1*) were immunoprecipitated above background levels, both of which have ≥ 1 CPEs in the 3’UTR, all low-TE transcripts with CPEs in the 3’UTR were efficiently recovered in the CPEB1-IP pellet.

### Binding of CPEB1 to CPE recruits a repressive complex in GV-arrested oocytes

To elucidate the mechanisms controlling translation during prophase I arrest, we focused on members of the oocyte-secreted protein (OOSP) cluster (*Oosp1*, *2*, and *3)* (Paillisson et al., 2005). *Oosp1* (red) and *Oosp3* (black) are translationally repressed during prophase I arrest and activated after meiotic resumption, while *Oosp2* is highly translated in GV-arrested oocytes and its translation becomes repressed during meiotic maturation (Fig. 5a). While 60% of *Oosp2* transcripts have poly(A) tails with ≥ 80 nts, *Oosp1* (21%) and *Oosp2* (6%) do not show this bias (Fig. 5b). The 3’UTRs of *Oosp1* and *Oosp3* have two and four putative CPEs upstream of the PAS, respectively, while *Oosp2* has no obvious CPE.

**Fig. 5.**
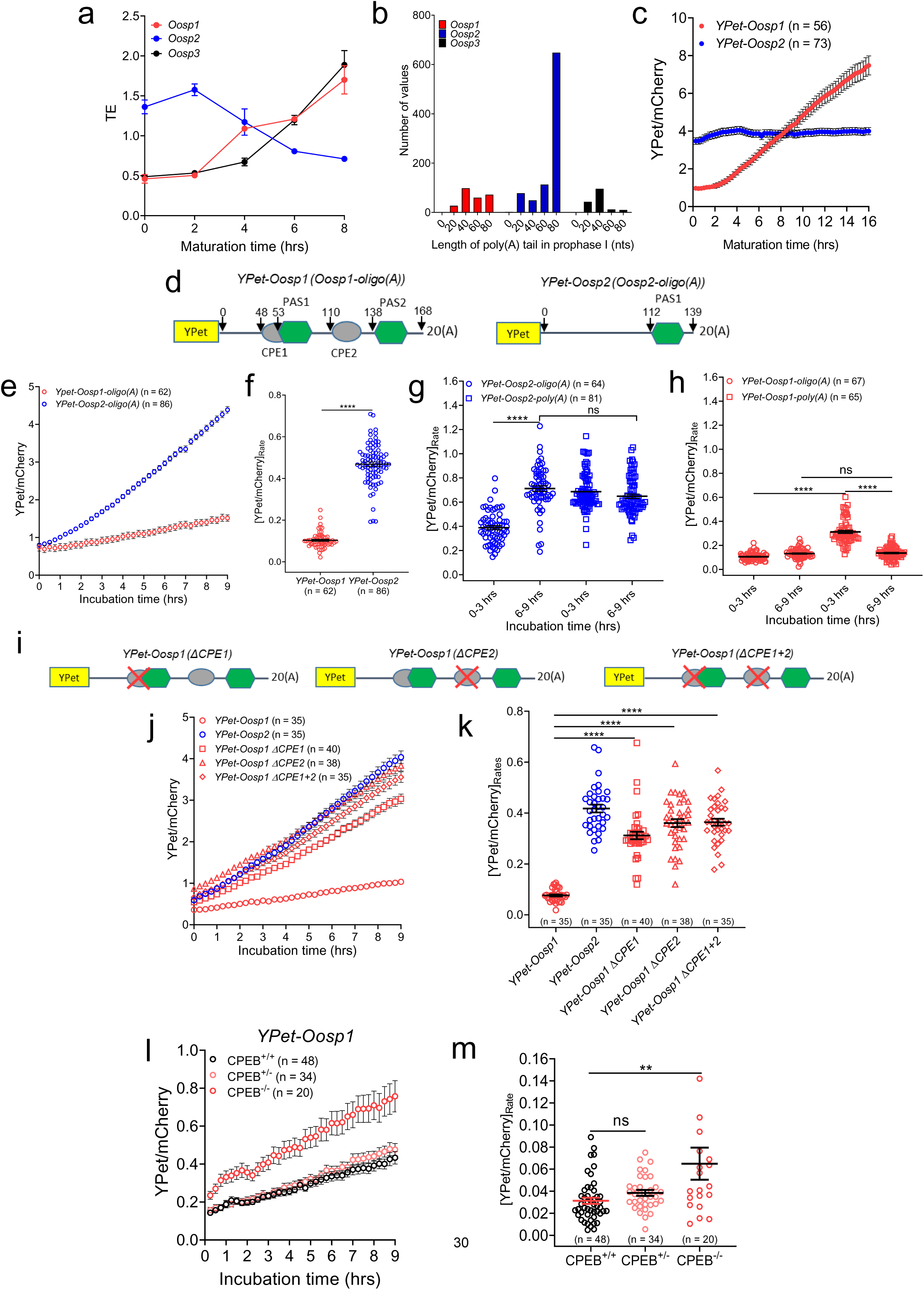
The presence of CPEs in the 3’UTR is associated with translational repression in GV-arrested oocytes. **a)** TE values of members of the *Oosp* cluster during meiosis. The dubplicate average and range of TEs are reported. **b)** Polyadenylation state of members of the *Oosp* cluster in GV-arrested oocytes. Data were from a published TAIL-Seq study(Morgan et al., 2017). **c)** Accumulation of YPet reporters for *Oosp1* and *Oosp2* 3’UTRs during meiotic maturation. GV-arrested oocytes were collected and microinjected with oligoadenylated *YPet*-*Oosp1* (red) or *YPet*-*Oosp2* (blue) mRNA along with polyadenylated *mCherry* mRNA. Oocytes were allowed to recover for 16 hrs after microinjection, released from cilostamide block, and imaged for 16 hrs with a sampling frequency of 15 mins. Each point is the mean ± SEM of individual oocyte traces obtained in three separate experiments. The total number of oocytes analyzed is in parentheses. **d)** YPet reporters for *Oosp1* and *Oosp2* 3’UTRs. 3’ UTRs expressed in the oocytes were cloned downstream of the YPet ORF (yellow box). CPEs (grey ovals) and PASes (green hexagons) are shown along with relevant nucleotide positions relative to the start of the 3’UTR. **e)** Accumulation of *Oosp1* and *Oosp2* 3’UTR YPet reporters in GV-arrested oocytes. GV-arrested oocytes were collected and microinjected with oligoadenylated *YPet*-*Oosp1* (red) or *YPet*-*Oosp2* (blue) mRNA along with polyadenylated *mCherry* mRNA. Oocytes were allowed to recover for 2.5 hrs after microinjection, maintained in prophase I, and imaged for 9 hrs with a sampling frequency of 15 mins. Each point is the mean ± SEM of individual oocyte traces obtained in three separate experiments. The total number of oocytes analyzed is in parentheses. **f)** Translation rates of the *Oosp1* and *Oosp2* YPet reporters in GV-arrested oocytes. The translation rate for each oocyte was calculated by linear regression of the reporter data (Fig. 5e) between 6 and 9 hrs. Mean ± SEM is reported. Statistical significance was evaluated by Mann Whitney test; ****: p < 0.0001. **g)** Translation rates of oligoadenylated or polyadenylated *Oosp2* 3’UTR YPet reporter in GV-arrested oocytes. GV-arrested oocytes were collected and microinjected with either *YPet-Oosp2-oligo(A)* or *YPet-Oosp2-poly(A)* mRNA along with polyadenylated *mCherry* mRNA. Experimental conditions were as described in Fig. 5e. The translation rate for each oocyte was calculated by linear regression of the reporter data (Supplemental Fig. 4c) between 0 and 3 hrs or 6 and 9 hrs. The data were collected from two independent experiments and the total number of oocytes analyzed and mean ± SEM are reported. Statistical significance was evaluated by Kruskal-Wallis test; ****: p < 0.0001 and ns: not significant. **h)** Translation rates of oligoadenylated and polyadenylated *Oosp1* YPet reporters in GV-arrested oocytes. Experimental conditions were as described in Fig. 5e. The data were collected from two independent experiments (Supplemental Fig. 4d) and the total number of oocytes analyzed and mean ± SEM are reported. Statistical significance was evaluated by Kruskal-Wallis test; ****: p < 0.0001 and ns: not significant. **i)** Mutations of CPE(s) in the *Oosp1* YPet reporter. We designated the proximal site as CPE1 and the distal as CPE2. CPE1 (TTTTAAATaaa) was mutated to ‘CGACAAATaaa,’, preserving the downstream, overlapping PAS, while CPE2 (TTTTAAT) was mutated to ‘CGACTCC’ as previously described(Yang et al., 2017). **j)** Accumulation of wild type *Oosp1*, wild type *Oosp2*, and mutant *Oosp1* reporters in GV-arrested oocytes. GV-arrested oocytes were collected and microinjected with oligoadenylated *YPet*-*Oosp1* (red circle), *YPet*-*Oosp2* (blue circle), *YPet*-*Oosp1(ΔCPE1)* (red square), *YPet*-*Oosp1(ΔCPE2)* (red triangle), or *YPet*-*Oosp1(ΔCPE1+2)* (red diamond) mRNA along with polyadenylated *mCherry* mRNA. Experimental conditions were as described in Fig. 5e. Each point is the mean ± SEM of individual oocyte traces obtained in two separate experiments. The total number of oocytes analyzed is in parentheses. **k)** Translation rates of wild type *Oosp1*, wild type *Oosp2*, and mutant *Oosp1* reporters in GV-arrested oocytes. The translation rate for each oocyte was calculated by linear regression of the reporter data (Fig. 5j) between 6 and 9 hrs. Mean ± SEM is reported. Statistical significance was evaluated by Kruskal-Wallis test; ****: p < 0.0001 and ns: not significant. **l)** Accumulation of *YPet-Oosp1* in GV-arrested CPEB1^+/+^, CPEB1^+/-^, and CPEB1^-/-^ oocytes. Oocytes were collected from hormone-primed wild type, *Zp3-Cre^T^ Cpeb1^F/+^*, and *Zp3-Cre^T^ Cpeb1^F/F^* mice. Experimental conditions were as described in Fig. 5e. Each point is the mean ± SEM of individual oocyte traces obtained in two separate experiments. The total number of oocytes analyzed is in parentheses. **m)** Translation rates of *YPet-Oosp1* in GV-arrested CPEB1^+/+^, CPEB1^+/-^, and CPEB1^-/-^ oocytes. The translation rate for each oocyte was calculated by linear regression of the reporter data (Fig. 5l) between 6 and 9 hrs. Mean ± SEM is reported. Statistical significance was evaluated by Kruskal-Wallis test; **: p = 0.0043 and ns: not significant.

Either *YPet-Oosp1* or *YPet-Oosp2* mRNAs (Fig. 5c and d) were injected into oocytes along with polyadenylated *mCherry* mRNA and the translation of YPet was monitored *via* quantitative live cell imaging (Supplementary Fig. 4a). The patterns of YPet accumulation for the *Oosp1* and *Oosp2* reporters recapitulate the translation patterns of the mRNAs during meiotic maturation as observed in the RNA-Seq data (Fig. 5c). During prophase I arrest, *YPet-Oosp1-oligo(A)* was translated at a significantly lower rate than *YPet-Oosp2-oligo(A)* (Fig. 5e and 5f). Notably, the translation rate of *YPet-Oosp2-oligo(A)* significantly increased during incubation, whereas translation rate of the polyadenylated reporter (*YPet-Oosp2-poly(A)*) was initially high and remained steady (Fig. 5g and Supplementary Fig. 4b and c). When polyadenylation of the *Oosp1* reporter was forced, the translation rate was initially high but decreased to levels comparable to those of the oligoadenylated reporter (Fig. 5h and Supplementary Fig. 4b and d). Therefore, regardless of its initial adenylation state, a reporter will eventually be translated at a rate dictated by the 3’UTR (Supplementary Fig. 4e).

In *YPet-Oosp1*, single as well as combined mutations of CPE1 and CPE2 in the 3’UTR (Fig. 5i) resulted in de-repressed translation to levels similar to those of *YPet-Oosp2* (Fig. 5j and k). These findings indicate a role of CPEs in the recruitment of an inhibitory complex. Using an oocyte-specific, CPEB1 loss-of-function model (Supplementary Fig. 5), we investigated the consequences of CPEB1 depletion on translation. While no significant difference in *Ypet-Oosp1* translation were detected between CPEB1^+/+^ and CPEB1^+/-^ oocytes, translation in CPEB1^-/-^ oocytes was significantly de-repressed (Fig. 5l and m). Of note, the effects of CPEB1 depletion on *Oosp1* de-repression were not as prominent as those achieved by mutating the CPEs in *Oosp1*. These results provide strong evidence that CPEB1 binding to CPEs is necessary for translation repression during prophase I arrest. However, regardless of the distance from the PAS, addition of a CPE into the 3’UTR of *Oosp2* was not sufficient to repress its translation rates (Supplementary Fig. 6).

### Translation repression during meiotic maturation is dependent on mRNA deadenylation and is dissociated from destabilization

From prophase I to late MI, ribosome loading for 1722 transcripts is decreased (DOWN, FDR ≤ 0.05). Translation repression is observed at as early as pro-metaphase I, but also occurs later on during meiosis (Fig. 1b and Supplementary Fig. 7 and 8). For 92% of DOWN transcripts, repression does not coincide with message destabilization, indicating that these two processes are mechanistically decoupled. Several DOWN candidates with stable mRNA levels were chosen for further investigation, including *Oosp2* and mRNAs that code for components of the *zona pellucida* (*Zp1*, *Zp2*, and *Zp3*) and the chromosome condensin complex (*Smc4*) (Fig. 6a). RT-qPCR confirmed the stability of these mRNAs, with all levels being constant up to 8 hrs and most up to 16 hrs of meiotic maturation (MII) (Fig. 6b). Poly(A) tail length (PAT) assays documents that *Smc4* and *Zp2* were polyadenylated in prophase I (Supplementary Fig. 9a) and, by 2 hrs, their poly(A) tails were significantly shortened (Fig. 6c).

**Fig. 6.**
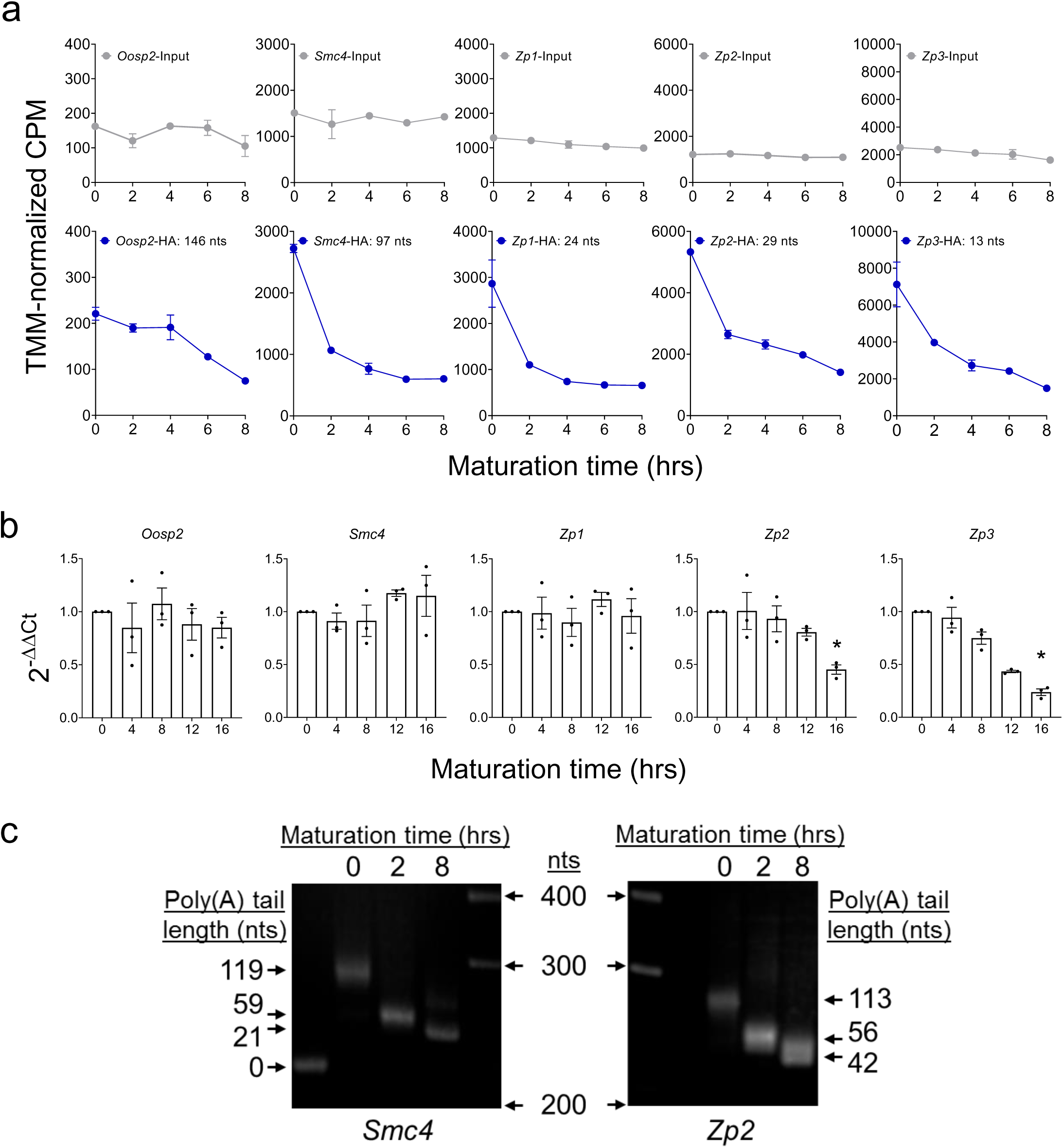
Translational repression during oocyte reentry into the cell cycle is dissociated from destabilization and requires de-adenylation. **a)** Time course of ribosome loading onto repressed candidate mRNAs (DOWN) during meiotic maturation. Values are from our RiboTag/RNA-Seq dataset and the mean and range of duplicate biological replicates are plotted. **b)** Translational repression of endogenous mRNAs is dissociated from destabilization. Oocytes were matured *in vitro* up to MII and samples were collected at different times during maturation. RNA was extracted from the oocytes, reverse transcribed, and used for RT-qPCR. *Bcl2l10* was used as a reference gene as its levels are known to be stable during this time. Data are represented as fold changes in mRNA levels as compared to 0 hrs. Three biological replicates of 30 oocytes per time point were used and RT-qPCR reactions were run in triplicate. The bars represent the mean ± SEM of three experiments. Statistical significance was evaluated by Friedman tests; *: p < 0.05. **c)** Translational repression of endogenous *Smc4* and *Zp2* is associated with message de-adenylation. Oocytes were either maintained in prophase I arrest (0 hrs) or allowed to mature for 2 or 8 hrs. At the end of the incubation, RNA was extracted and used for PAT assays with anchored oligo-dT primers. A representative experiment of the three performed is reported.

### A CPE in close proximity of the PAS is required to maintain translation during meiotic maturation

The *YPet-Zp2* reporter was translated at relatively high and steady rates during prophase I and translation was repressed shortly after GVBD (Fig. 7a and b), in agreement with the RNA-Seq data (Fig. 6a) and PAT assay (Fig. 6c). Similar results were obtained with *YPet-Smc4* (Supplementary Fig. 9b and c) and *Ypet-Oosp2* (see below). Oocytes released from cilostamide block and simultaneously treated with the CDK1 inhibitor dinaciclib did not show differences in *YPet-Zp2* translation rates when compared to oocytes maintained in cilostamide, suggesting that CDK1 activation and GVBD are required for translation repression. PKA inhibitor treatment (Rp-cAMPS), used to block cAMP signaling, again resulted in no repression, indicating that cAMP is not a signal involved in translation repression (Fig. 7c). However, treatment of oocytes with dinaciclib after GVBD (2 hrs) resulted in decreased translation repression as compared to control oocytes. Therefore, CDK1 activation and GVBD are events necessary to trigger translational repression of *Zp2* upon meiotic resumption.

**Fig. 7.**
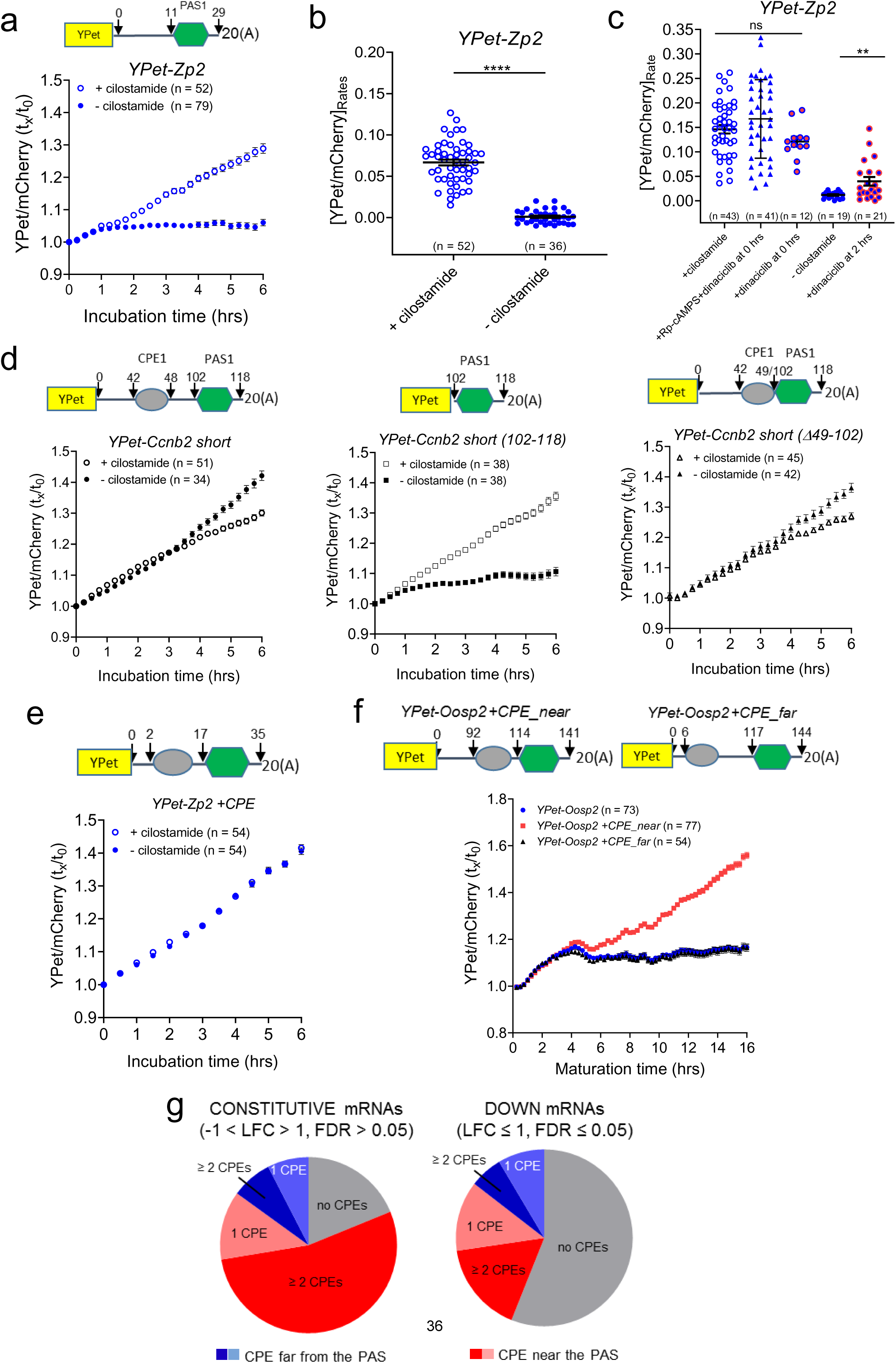
Translational repression during meiotic maturation is recapitulated by the 3’UTR of DOWN mRNAs, requires CDK1 activation, and is prevented by the presence of a CPE in close proximity to the PAS. **a)** The 3’ UTR of *Zp2* (high-TE in prophase I and DOWN transcript) recapitulates the rapid translation repression post-GVBD. Oocytes were injected with an oligoadenylated *YPet-Zp2* mRNA together with a polyadenylated *mCherry* mRNA. Oocytes were then either maintained in prophase I arrest with cilostamide treatment (empty circles) or allowed to mature (solid circles) and imaged for 10 hrs with a sampling frequency of 30 mins. Data are reported as the fold change of the YPet/mCherry ratios as compared to 0 hrs. Each point is the mean ± SEM of individual oocyte traces obtained in two separate experiments. The total number of oocytes analyzed is in parentheses. **b)** Translation rates of the *YPet-Zp2* reporter in GV-arrested or maturing oocytes. The translation rate for each oocyte was calculated by linear regression of the reporter data (Fig. 7a) between 3 and 6 hrs. Mean ± SEM are reported. Statistical significance was evaluated by unpaired, two-tailed t-test; ****: p < 0.0001. **c)** Translation repression of the *YPet-Zp2* reporter during meiosis resumption requires GVBD and CDK1 activation but not PKA activity. After microinjection of the *YPet-Zp2* reporter, oocytes were released in cilostamide-free medium and incubated with a CDK1 inhibitor (5 μM dinaciclib) or a combination of CDK1 and PKA inhibitors (Rp-cAMPS) from the time of release (0 hrs). The translation rate for each oocyte was calculated by linear regression of the reporter data between 3 and 6 hrs. In another group, dinaciclib was added after GVBD at 2 hrs into incubation. Statistical significance was evaluated by unpaired, two-tailed t-tests; ns: not significant; **: p = 0.0053. **d)** Deletion mutagenesis of the *Ccnb2* 3’UTR. GV-arrested oocytes were collected, microinjected with oligoadenylated *YPet-CcnB2 short* or reporters fused to the *Ccnb2* 3’UTRs with progressive deletions along with a polyadenylated *mCherry* reporter. Sixteen hrs after microinjection, oocytes were either maintained in prophase I arrest with cilostamide (empty circles) or allowed to mature (solid circles) and imaged for 6 hrs with a sampling frequency of 15 mins. Data are reported as the fold change of the YPet/mCherry ratios as compared to 0 hrs. Each point is the mean ± SEM of individual oocyte traces obtained in two separate experiments. The total number of oocytes analyzed is in parentheses. **e)** Insertion of a CPE in the 3’ UTR of *Zp2* (DOWN), prevents repression during meiotic maturation. GV-arrested oocytes were collected, microinjected with oligoadenylated *YPet-Zp2 +CPE* mRNA together with a polyadenylated *mCherry* mRNA. Experimental conditions were as described in Fig. 7a. Data are reported as the fold change of the YPet/mCherry ratios as compared to 0 hrs. Each point is the mean ± SEM of individual oocyte traces obtained in two separate experiments. The total number of oocytes analyzed is in parentheses. **f)** Insertion of a CPE in close proximity of the PAS in the *Oosp2* 3’UTR prevents translational repression during meiotic maturation. GV-arrested oocytes were collected, microinjected with oligoadenylated *YPet-Oosp2* or a reporter with a CPE inserted in the *Oosp2* 3’UTR along with a polyadenylated *mCherry* reporter. Oocytes were incubated for 16 hrs after microinjection, allowed to mature, and imaged for 16 hrs with a sampling frequency of 15 mins. Data are reported as the fold change of the YPet/mCherry ratios as compared to 0 hrs. Each point is the mean ± SEM of individual oocyte traces obtained in three separate experiments. The total number of oocytes analyzed is in parentheses. **g)** Detailed analysis of the relationship between translation patterns during meiotic resumption and the presence of CPEs in the 3’UTR. Pie charts report the percentage of CONSTUTIVE or DOWN mRNAs in GV-arrested oocytes that have or lack CPEs in the 3’UTR.

To elucidate the mechanisms of constitutive or repressed translation during meiotic maturation, we monitored the translation of the *CcnB2* reporter, which is constitutively translated before and after GVBD. Progressive deletions of the *Ccnb2 short* 3’UTR revealed that a reporter retaining only the PAS sequence (*YPet-CcnB2 short (102-118)*) was translated like a prototypical DOWN gene; it was highly translated during prophase I and translation became repressed after GVBD (Fig. 7d). Presence of the first 42 nts of the 3’UTR did not significantly affect the translation pattern (Supplementary Fig. 10a). If the 3’UTR included a CPE (*YPet-Ccnb2 short (Δ49-102)*, translation of the reporter was no longer repressed post-GVBD (Fig. 7d) and resembled the pattern of a prototypical CONSTITUTIVE mRNA. Therefore, the presence a CPE is critical for an mRNA to evade translation repression during meiotic maturation. This was confirmed by a gain-of-function experiment using a repressed mRNA, where insertion of a CPE was sufficient to maintain the high, prophase I translation rate of *YFP-Zp2* after meiotic resumption (Fig. 7e).

We tested whether CPE position in relation to the PAS is important for the maintenance of translation post-GVBD by using the *Oosp2* 3’UTR. When a CPE was added 22 nts upstream of the PAS, reporter translation was no longer repressed upon meiotic resumption, but maintained at the constant rate as in prophase I (Fig. 7f and Supplementary Fig. 10b). However, if the same CPE was added 111 nts upstream of the PAS, translation was still repressed post-GVBD and the pattern did not differ from that of *YPet-Oosp2* (Fig. 7f and Supplementary Fig. 10b). Therefore, during meiotic maturation, inclusion of a CPE proximal to the PAS in the 3’UTR of a translationally repressed mRNA (gain-of function) changed its translation pattern to that of a constitutively translated mRNA. Conversely, removal of a CPE proximal to the PAS from the 3’UTR of a constitutively translated mRNA (loss-of-function) transformed its translation pattern to that of a repressed mRNA.

Genome-wide analysis shows that 82% of CONSTITUTIVE mRNAs have at ≥ 1 CPEs in the 3’UTR, while this is true for only 47% of DOWN mRNAs (Fig. 7g). Further analysis of these classes reveals a bias towards the presence of CPEs within 50 nts upstream of the PAS in constitutively translated mRNAs (Supplementary Fig. 11).

### Activation of translation during meiotic maturation is dependent on GVBD and, in part, activation of CDK1

Concurrent with translation repression, progression through meiosis is associated with significant increases in translation of 1537 maternal mRNAs (UP, FDR ≤ 0.05) (Supplementary Fig. 1e). These UP mRNAs show both early and late increased translation (Supplementary Fig. 12). To investigate the mechanisms underlying this activation, we chose candidates with some of the highest fold changes in ribosome loading from prophase I to late MI, *Tcl1*, *Oosp1*, *Obox5*, *Ccnb1*, and *Ewsr1* (Fig. 8a). RiboTag/RT-qPCR confirmed the translation pattern of these UP transcripts (Fig. 8b). To investigate the link between cell cycle and the translation program during meiosis, we used a YPet reporter fused with the *Ccnb1* 3’UTR, a transcript whose translation activation after GVBD has been shown to be CDK1-dependent (Han et al., 2017). When CDK1 activity was inhibited immediately after release from PDE inhibition, GVBD did not occur and translation was maintained at levels prior to cilostamide release (Fig. 8c). Oocytes treated with dinaciclib after GVBD (2 hrs) eventually regained a nuclear membrane, indicating effective CDK1 inhibition, and translation of *Ypet-Ccnb1* was reduced, but not abolished. This experiment was also performed using *YPet-Ewsr1*, *YPet-Oosp1*, and *YPet-Mos* (Supplementary Fig. 13). While dinaciclib treatment of GV-arrested oocytes completely inhibited translation of all the reporters, CDK1 inhibition after GVBD only decreased the translation of *YPet-Ccnb1 and Ypet-Ewsr1*; the translation of *YPet-Oosp1* and *YPet-Mos* was unaffected (Fig. 8d). Therefore, early CDK1 activity responsible for GVBD is required for translation activation of these candidates, while CDK1 activity in MI is only partially responsible for subsequent increases in translation of *Ccnb1* and *Ewsr1*. The variable effects of dinaciclib may be due to subtle differences in the timing and mechanism’s of translation activation.

**Fig. 8.**
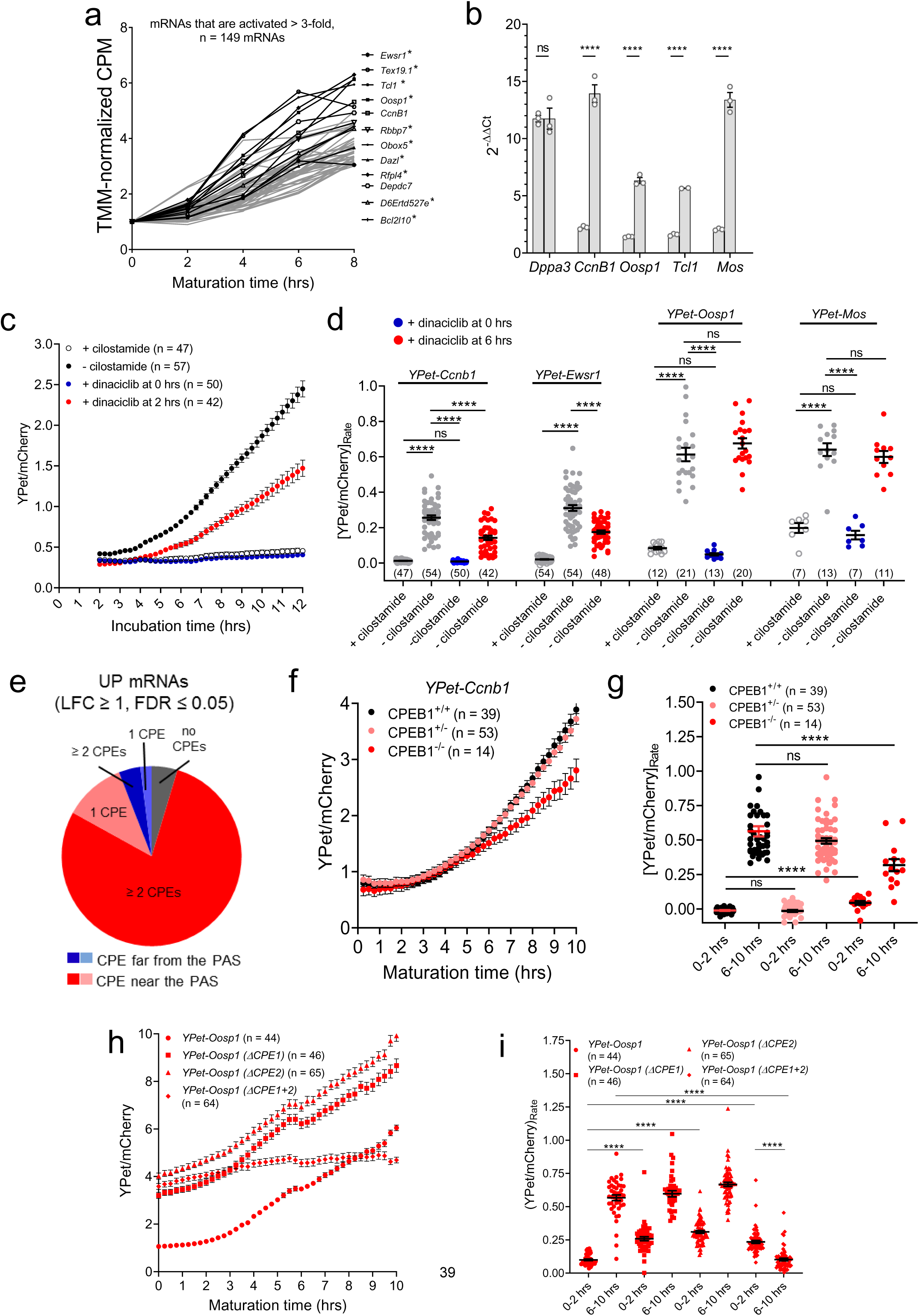
CPEB binding to mRNAs activated during maturation is necessary, but not sufficient, for full translational activation. **a)** Pattern of ribosome loading onto UP mRNAs during meiotic maturation. mRNAs whose translation increased by at least three-fold from prophase I to late MI in our RiboTag/RNA-Seq dataset are shown. Traces of the 149 mRNAs with the highest activation are in grey and transcripts recovered in the pellet of RNA-IP/RT-qPCR with CPEB1 are in black. * denotes transcripts that are also immunoprecipitated by DAZL antibodies (data under review). **b)** RiboTag-IP/RT-qPCR validation of ribosome loading for selected UP candidates. *Zp3-Cre^T^ RiboTag^F/F^* mice were hormone primed and the oocytes isolated. Oocytes were either maintained in prophase I or matured *in vitro* for 8 hrs and collected for downstream RiboTag-IP/RT-qPCR analysis. We quantified several candidates with some of the greatest fold changes in ribosome loading from prophase to late MI. *Dppa3* was used as a reference gene as it is known to be constitutively translated during this time. Data are represented as fold changes in message levels as compared to 0 hrs. Three biological replicates of 200 oocytes per time point were used and RT-qPCR reactions were run in triplicate. The bars represent the mean ± SEM of three experiments. Statistical significance was evaluated by unpaired, two-tailed t-tests; ****: p < 0.0001. **c)** The effect of CDK1 inhibition on the translation of *CcnB1* (UP). GV-arrested oocytes were collected and microinjected with oligoadenylated *YPet*-*CcnB1* 3’UTR mRNA along with polyadenylated *mCherry* mRNA. Oocytes were incubated for 16 hrs then two groups of oocytes were maintained in prophase I arrest with either cilostamide (empty, black circle) or dinaciclib without cilostamide (blue circle). Another two groups of oocytes were either matured without (solid, black circle) or with dinaciclib added at 2 hrs after release (red circle). Imaging started 2 hrs after cilostamide release and lasted for 10 hrs with a sampling frequency of 15 mins. Each point is the mean ± SEM of individual oocyte traces obtained in three separate experiments. The total number of oocytes analyzed is in parentheses. **d)** Translation rates of *YPet*-*CcnB1* and *YPet-Ewsr1* are affected by CDK1 inhibition during meiotic maturation. The translation rate for each oocyte was calculated by linear regression of the reporter data (Fig. 8c and Supplementary Fig. 13a-c) between 8 and 12 hrs. Mean ± SEM is reported. Statistical significance was evaluated by Kruskal-Wallis test; ns: not significant; ****: p < 0.0001. **e)** Detailed analysis of the relationship between mRNAs that are translationally activated during meiotic resumption and the presence of CPEs in the 3’UTR. Pie charts report the percentage of UP mRNAs in GV-arrested oocytes that have or lack CPEs in the 3’UTR. **f)** CPEB1 is required for efficient translational activation of *CcnB1*. CPEB1^+/+^ (black), CPEB1^+/-^ (light red), and CPEB1^-/-^ (red) oocytes were collected, maintained in prophase I arrest, and microinjected with oligoadenylated *YPet*-*CcnB1* mRNA along with polyadenylated *mCherry* mRNA. After 2.5 hrs incubation, oocytes were matured and imaged for 10 hrs with a sampling frequency of 15 mins. Each point is the mean ± SEM of individual oocyte traces obtained in two separate experiments. The total number of oocytes analyzed is in parentheses. **g)** Translation rates of the *YPet*-*CcnB1* reporter during oocyte maturation in CPEB1^+/+^, CPEB1^+/-^, and CPEB1^-/-^ oocytes. The translation rate for each oocyte was calculated by linear regression of the reporter data (Fig. 8f) between 0 and 2 hrs or 6 and 10 hrs. Mean ± SEM is reported. Statistical significance was evaluated by Kruskal-Wallis test; ns: not significant; ****: p < 0.0001. **h)** Accumulation of wild type *Oosp1* and mutant *Oosp1* YPet reporters during meiotic maturation. GV-arrested oocytes were collected and microinjected with oligoadenylated *YPet*-*Oosp1* (circle), *YPet*-*Oosp1(ΔCPE1)* (square), *YPet*-*Oosp1(ΔCPE2)* (triangle), or *YPet*-*Oosp1(ΔCPE1+2)* (diamond) mRNA along with polyadenylated *mCherry* mRNA. After 16 hrs of recovery after microinjection, oocytes were allowed to mature, and imaged for 10 hrs with a sampling frequency of 15 mins. Each point is the mean ± SEM of individual oocyte traces obtained in two separate experiments. The total number of oocytes analyzed is in parentheses. **i)** Translation rates of wild type *Oosp1* and mutant *Oosp1* YPet reporters during meiotic maturation. The translation rate for each oocyte was calculated by linear regression of the reporter data (Fig. 8f) between 0 and 2 hrs or 6 and 10 hrs (post-GVBD). Mean ± SEM is reported. Statistical significance was evaluated by Kruskal-Wallis test; ns: not significant; ****: p < 0.0001.

Regulation of mRNA translation by CDK1 is thought to be mediated by phosphorylation of CPEB1 (Ballantyne et al., 1997; Han et al., 2017). Genome-wide analysis reveals that 95% of UP transcripts have ≥ 1 CPEs in the 3’UTR (Fig. 8e). Using CPEB1^-/-^ oocytes, we investigated the role of CPEB1 in the regulation of *Ccnb1* translation. CPEB1^-/-^ oocytes showed significantly decreased, but not abolished, translation rates as compared to wild type oocytes (Fig. 8f and g). Moreover, depletion of CPEB resulted in higher translation rates prior to GVBD (Fig. 8g), confirming the role of this RBP in translation repression during prophase I. Single mutations of CPE1 and CPE2 in *Oosp1* resulted in a significant increase in initial reporter translation, but there were no effects on translation activation post-GVBD (Fig. 8h and i). This indicates that both CPEs are necessary for translation repression during prophase I, but only one CPE is sufficient for translation activation. Mutation of both CPEs did not further de-repress translation before GVBD, but completely abolished translation activation post-GVBD (Fig. 8h and i), suggesting that translation activation is not simply due to de-repression and that the two processes are dissociated.

## Discussion

Female gamete development is driven by transcription of not only maternal mRNAs essential for the expansive growth taking place during this phase, but also of mRNAs needed for the synthesis of proteins used later on during meiotic progression and embryo development. To accomplish this developmental program, not all mRNAs are translated immediately after synthesis. Some are, instead, stored for future use during later stages of development and their translation is repressed until then. In this study, we used a genome-wide approach to demonstrate that translation of housekeeping and instructive mRNAs in quiescent GV-arrested oocytes is due only in part to low complexity 3’UTRs and therefore is not the default pathway. Robust translation of 84% of high-TE mRNAs is associated with 3’ UTRs that are > 200 nts and include putative binding elements for known RBPs. Conversely, translation repression in quiescent oocytes is dependent on the assembly of complexes that promote deadenylation; our data suggest that this complex is nucleated by CPEB1. CPEB1 is not sufficient, though, as repression is mitigated, not abolished, in CPEB1^-/-^ oocytes. Either compensatory mechanisms or alternative repressive networks are functional in these GV-arrested oocytes. As the oocyte resumes meiosis, a switch in the translation program occurs at around the time of germinal vesicle breakdown. Messenger RNAs translated at high levels during growth become deadenylated and repressed in a manner dependent on the properties of the 3’UTR, but total levels remain stable for the majority of these mRNAs. Conversely, translation of mRNAs that were repressed in quiescent oocytes becomes progressively activated through MI and MII.

The timing of translational repression is unique for each mRNA. In frog oocytes, it is proposed that the deadenylase PARN is released at nuclear envelope breakdown, inducing the default process of deadenylation (Barnard et al., 2004; Mendez et al., 2000a; Mendez et al., 2000b). This mechanism may apply to only a minority of mRNAs in mouse oocytes, as 77% of the maternal mRNA analyzed are translationally repressed well after GVBD, during MI or later. It should be noted that, in some cases, translation repression coincides with inactivation of the encoded protein. For example, CPEB1 protein degradation initiates at four to six hours post-meiotic resumption (Han et al., 2017; Mendez et al., 2002). Our RNA-Seq data indicate that ribosome association with *Cpeb1* mRNAs begins to decrease around this time. Translation of *Cdk1* similarly decreases at the MI-to-anaphase transition, when M-Phase-promoting factor (MPF) is inactivated because of CCNB1 degradation (Evans et al., 1983). Thus, changes in message translation likely cooperates with protein turnover to regulate functions during meiosis.

Our genome-wide analysis of translation demonstrates that the environment of stable maternal mRNAs present during oocyte growth extends well into the late stages meiosis. Here, we have identified a first wave of mRNA destabilization at the MI-to-anaphase transition that affects a subset of mRNAs; most messages that are translationally activated along with a subset of translationally repressed mRNAs continue to remain stable well into MII. Eventually, these maternal messages will be eliminated later on in the zygote and in the embryo at the time of zygote genome activation in two successive waves of degradation (Yartseva and Giraldez, 2015). Recently, mRNA methylation has emerged as a key regulator of mRNA stability and the m6A reader YTHDF2 has been implicated in message destabilization in the oocyte (Ivanova et al., 2017; Wang et al., 2014). However, we found minimal overlap between mRNAs destabilized in MI and those stabilized by YTHDF2 loss-of-function. Thus, the code controlling timed destabilization remains poorly understood. However, a CCR4/CNOT complex that includes CNOT7/8 and BTG4 is responsible for destabilization of a subset of repressed mRNAs(Liu et al., 2016; Pasternak et al., 2016; Yu et al., 2016). Our RiboTag/RNA-Seq data do indicate that the machinery required for mRNA destabilization/degradation is synthesized late during oocyte maturation due to delayed translation of mRNAs such as *Btg4*, *Cnot7*, and *Dcp1a*.

CPEB1 is considered a master regulator of translation during oocyte meiosis. Our findings are consistent with this concept, as 95% of mRNAs significantly activated from prophase I to late MI have at least one CPE in the 3’UTR and all translationally activated candidates we tested interact with CPEB1. Depletion of CPEB1 in the oocyte did not completely abolish translation activation, pointing to the presence of other RBPs with similar function to CPEB1 or other compensatory mechanisms. Moreover, we have characterized an additional function of this RBP during oocyte maturation: CPEB1 is required for maintaining translation of thousands of maternal mRNAs, preventing their deadenylation. Our genome-wide data provide evidence that a CPE in close proximity of the PAS is required for this widespread translation of the majority of maternal mRNAs. Studies in frog oocytes have revealed the presence of a combinatorial code of CPEs enforcing repression and early and late activation(Pique et al., 2008). Our global analysis documents that CPE elements cluster in the vicinity of the PAS (< 100 nts), and that both repression and activation are associated with the presence of more than one CPE element in the 3’UTR. Using *Oosp1* mRNA as prototypic mRNA repressed in prophase and activated in MI, we demonstrated that two CPEs are required for repression, but one CPE is sufficient for activation. Although some of the features of the combinatorial code described in *Xenopus* were confirmed, more complex rules govern the functions of CPEs in oocyte translation.

We demonstrate that the pattern of translation switches at the time CDK1 is activated in the oocyte. If CDK1 activity is inhibited, no activation or repression of translation takes place and GVBD, however, is also blocked. Under our experimental conditions and if CDK1 inhibition is delayed after GVBD, translation of some mRNAs including *CcnB1* and *Ewsr1* was partially inhibited whereas that of *Mos* and *Oosp1* was not significantly affected. A possible explanation of this differential sensitivity to CDK1 is the presence of distinct mechanisms of translational activation downstream of CDK1. Differences in translation mechanisms for *Mos* and *Ccnb1* have been demonstrated in frog oocytes(Mendez et al., 2002; Sheets et al., 1994). However, given the very rapid re-entry in meiosis of mouse oocytes, we are unable to dissect possible CDK1-independent mechanisms of translation activation.

In summary, our genome-wide approach provides a novel perspective on the dynamics of the translational program in mouse oocytes. Since the combinatorial code described in frog oocytes(Pique et al., 2008) is applicable only to a subset of mammalian mRNAs, more complex mechanisms are functioning in mammalian oocytes. Additionally, to be defined is whether a CPEB-mediated translation program is required during oocyte growth as short, low complexity 3’UTRs are associated with only a portion of the mRNA translated during growth. Finally, differences in how the translation program is executed between mouse and human must exist and should to be explored to gain a better understanding of the human oocyte-to-zygote transition and female fertility.

## Acknowledgments

We are indebted to John C. Wright (University of California, Berkeley) for assistance with bioinformatics analyses, as well as Elena Gochez (San Francisco VA Health Care System) and Emily Miller (Stanford University) for technical assistance. We thank Dr. Raúl Méndez and Dr. Gonzalo Fernandez-Miranda (The Barcelona Institute of Science and Technology) for sharing the CPEB1-targeted mice. X.G.L. was supported by 5T32HD7263-35 and E.M.D. was supported by the Lalor Foundation postdoctoral fellowship. This work was supported by National Institutes of Health: NIH R01 GM116926 and P50 HD055764.

## Author Contributions

Conceptualization: X.G.L., E.M.D., and M.C.; Methodology: X.G.L., E.M.D., and M.C.; Software: X.G.L.; Validation: X.G.L. and E.M.D.; Formal Analysis: X.G.L., E.M.D., and M.C.; Investigation: X.G.L., E.M.D. G.R., and C.Y.; Resources: M.C.; Data curation: X.G.L.; Writing – Original Draft: X.G.L., E.M.D., and M.C.; Writing – Review & Editing: X.G.L., E.M.D., G.R., C.Y., and M.C.; Visualization: X.G.L. and E.M.D.; Supervision: M.C.; Project administration: X.G.L., E.M.D., and M.C.; Funding acquisition: X.G.L., E.M.D., and M.C. X.G.L. and E.M.D. contributed equally to this study.

## Methods

### Animals

All experimental procedures involving mice were approved by the University of California, San Francisco Institutional Animal Care and Use Committee (Approval #AN101432). Animal care and use were performed according to relevant guidelines and regulations. All animals used were of the C57BL/6J inbred strain. C57BL/6-Zp3cre-Rpl22tm1.1Psam (*Zp3-Cre^T^ RiboTag*) mice were obtained from Jackson Laboratories and bred as previously described(Martins et al., 2016). CPEB1-targeted mice were a gift from Raúl Méndez and colleagues(Calderone et al., 2016) and bred in our laboratory.

### Oocyte isolation and culture

Three-week old female mice were injected with 5 I.U. PMSG to induce superovulation. Forty-four hrs after injection, the mice were euthanized and the ovaries dissected into media containing HEPES and 1 µM cilostamide (Millipore, 231085) (HC media). The antral follicles were punctured, allowing release of cumulus-oocyte complexes (COCs). Repeated aspiration through a glass pipette allowed for removal of the surrounding cumulus cells. Denuded oocytes were maintained at prophase I arrest in MEM Alpha (Gibco, 12561-056) supplemented with sodium pyruvate (Gibco, 11360070) and penicillin-streptomycin (Gibco, 15140122) in addition to 1 µM cilostamide (αC media) at 37°C under 5% CO_2_. If indicated, oocytes were transferred to cilostamide-free MEM Alpha (α media), allowed to mature, and collected at various time points. Where specified, oocytes were treated with 5 µM dinaciclib (Selleckchem, SCH727965) or 10 mM Rp-cAMPS.

### Immunofluorescence staining and confocal microscopy

Oocytes were collected at various time points and fixed in DPBS (GE Healthcare, SH30264.02) supplemented with 0.1% Triton X-100 (Sigma, X-100) and 2% formaldehyde (ThermoFisher, 28908) for 30 mins. After washing in blocking buffer (1x DPBS, 0.3% BSA, and 0.01% Tween), the oocytes were incubated in blocking buffer for 16 hrs and permeabilized for 15 mins in DPBS supplemented with 0.3% BSA and 0.1% Triton X-100. Samples were washed and then incubated for 1 hr with primary antibody diluted in blocking buffer. The antibodies used were: 1:100 β-tubulin (9F3) rabbit mAb (Cell Signaling Technology, 3623); 1:200 human antibody against centromere (ImmunoVision, HCT-0100). After another round of washing, samples were incubated for 1 hr with the corresponding secondary antibody, goat anti-rabbit IgG, Alexa Fluor 488 (ThermoFisher, A-11008) or 1:500 goat anti-human IgG, Alexa Fluor 568 (ThermoFisher, A-21090). Oocytes were washed again and then mounted with VECTASHIELD® Antifade Mounting Medium with DAPI (Vector, H-1200). All washes were done three times each round in blocking buffer for 10 mins each wash. Images were captured with a confocal Nikon C1SI equipped with X60 oil immersion lens and processed with ImageJ(Rueden et al., 2017).

### RiboTag-immunoprecipitation (RiboTag-IP)

Only *Zp3-Cre^T^ RiboTag^F/F^* female mice used for RiboTag-immunoprecipitation. Oocytes were collected in 5 µl 0.1% polyvinylpyrrolidone (PVP; Sigma, P0930) in 1x PBS (Invitrogen, AM9625), flash frozen in liquid nitrogen, and stored at −80°C.

The appropriate volume (50 µl per sample) of Dynabeads™ Protein G (Invitrogen, 10004D) was washed three times in 500 µl homogenization buffer (HB: 50 mM Tris HCl pH 7.4, 100 mM KCl, 12 mM MgCl_2_, and 1% NP-40) on a rotor at 4°C for 5 mins per wash. Two additional washes were performed with 500 µl sHB on a rotor at 4°C for 10 mins per wash. The final wash solution was removed and the beads were eluted in the original volume of HB supplemented (sHB) with 1mM DTT, 1x protease inhibitors, 200 units/ml RNaseOUT, 100 µg/ml cycloheximide, and 1mg/ml heparin. Samples were thawed, randomly pooled to yield a total of 200 oocytes per time point per replicate, and 300 µl of sHB was added to each pooled sample. To lyse the cells, samples were vortexed for 30 secs, flash frozen in liquid nitrogen, and thawed at room temperature (RT); this process was repeated twice. Finally, the homogenates were centrifuged for 10 mins at maximum speed and 4°C and the supernatants were transferred to new tubes. To pre-clear the samples, 20 µl washed beads was added to each supernatants and samples were incubated on a rotor at 4°C for 1 hr. A magnetic rack was used to remove the beads and 15 µl of each pre-cleared lysate was collected and added to 200 µl RLT buffer per sample (Qiagen, 74034) to serve as the input samples. The input samples were frozen and kept at −80°C until RNA extraction. Three µl (3 µg) anti-HA.11 epitope tag antibody (BioLegend, 901501) was added to each of the extracts and all samples were incubated on a rotor at 4°C for 4 hrs. Thirty µl washed beads was then added to the samples and incubated overnight on a rotor at 4°C. The beads (now bound by HA-tagged ribosomes and the associated mRNAs) were washed 5 times in 1 ml of urea wash buffer (uWB: 50 mM Tris HCl pH 7.4, 150 mM KCl, 12 mM MgCl_2_, 1% NP-40, 1x protease inhibitors, 1 mM DTT, 40 U RNaseOUT, 1 mg/ml heparin, 100 μg/ml cycloheximide, and 1 M urea) on a rotor at 4°C for 10 mins per wash. The beads were then pelleted via a magnetic rack and the uWB removed. Two-hundred and fifty µl RLT buffer was added to each sample and the samples were vortexed for 30 seconds. RNA extraction was performed following the Rneasy Plus Micro Kit protocol (Qiagen, 74034). Samples were eluted in 10 µl of RNase-free water and used downstream for RNA-Seq or RT-qPCR analysis.

### RNA-Seq

#### RNA sequencing

RNA samples were sent to the Gladstone Institutes Genomics Core for quality control using Bioanalyzer (Agilent) and cDNA library preparation. Samples were sequenced using the HiSeq 4000 system (Illumina).

#### Sequence quality assessment and trimming

The quality of the raw sequence data was checked via FASTQC. The sequence files were then trimmed using tools in Trimmomatic-0.36(Bolger et al., 2014). The following were removed: Illumina TruSeq3 single-ended adapter sequences, bases with a quality score lower than 3 at the start and end of a read, bases that had an average quality per base of below 15 calculated using a sliding window to average four bases, and any reads that were shorter than 36 bases. Input reads were single-ended and input qualities were ASCII characters equal to the Phred quality plus 33.

#### Mapping and counting reads

HiSat2(Kim et al., 2015) was used to build indexes from the Reference Consortium Mouse Build 38 (mm10) and to align sequence reads to the genome. The resulting .bam files were sorted and indexed with SAMtools(Li et al., 2009). Count files for each group were created with HTSeq using the Mouse GENCODE Gene set release M11. The input data were .bam files, the data were not from a strand-specific assay, and the feature type used was ‘gene.’

#### Differential expression (DE) analysis

The Bioconductor packages edgeR(Robinson et al., 2010) and limma(Ritchie et al., 2015) were used for statistical analyses. Only reads with greater or equal to 10 counts per million (CPM) in at least 2 samples were kept. Trimmed mean of M-values (TMM) normalization, which accounts for compositional differences among the libraries, was then performed on HA reads and input reads, separately. Using the raw counts, dispersion, and design matrix, the negative binomial generalized linear model was fitted for each gene. Finally, pairwise likelihood ratio tests for 2, 4, 6, and 8 hrs versus 0 hrs were conducted.

#### Gene ontology (GO) analysis

Gene lists were uploaded to DAVID 6.8(Huang et al., 2009a, b) and processed with the Functional Annotation Tool.

#### Analysis of 3’UTR sequences

The 3’UTR sequences of the genes of interest (including known mRNA isoforms) were downloaded using the Table Browser (UCSC Genes track) provided by GBShape(Chiu et al., 2015). The locations of putative PAS (AATAAA, ATTAAA, and AAGAAA)(Beaudoing et al., 2000) and CPE (TTTTAT, TTTTAAT, TTTTACT, TTTTAAAT, TTTTAAGT, and TTTTCAT)(Pique et al., 2008) sequences were determined via the Find Individual Motif Occurrences (FIMO) tool(Grant et al., 2011), which is part of the MEME Suite(Bailey et al., 2009); only exact matches were used for downstream analysis. Python scripts were written to calculate the distance of each CPE from each PAS for individual 3’UTRs.

#### Data visualization

ggplot2(Wickham, 2009) was used to create the multi-dimensional scaling (MDS) plots, which are used to visualize sample-to-sample distances, and translation activation and repression time course plots. All other graphs were plotted using GraphPad Prism 8.

### RNA immunoprecipitation (RNA-IP)

The appropriate volume (50 µl per sample) of Dynabeads™ Protein G (Invitrogen, 10004D) was washed twice in 250 ul incomplete lysis buffer (iLB: 15 mM Tris HCl pH 7.4, 75 mM NaCl, 5 mM MgCl_2_, 0.25% NP-40, 0.125 mM Na_3_VO_4_, and 5 mM β-glycerophosphate) on a rotor at 4°C for 5 mins. Two additional washes were performed with 250 µl complete LB (cLB: iLB supplemented with protease inhibitor, DTT, RNAseOUT, ribonucleoside vanadyl complex, and cycloheximide) on a rotor at 4°C for 5 mins per wash. The final wash solution was removed and the beads were eluted in the original volume of cLB. Oocytes were isolated as described and kept arrested at prophase I. Two hundred oocytes were collected in 5 µl 0.1% PVP in PBS, flash frozen in liquid nitrogen, and stored at −80°C. To homogenize the cells, 250 µl of cLB was added to the samples, samples were vortexed for 30 secs, and then incubated on ice for 10 mins. The homogenates were then centrifuged for 10 mins at maximum speed at 4°C and the supernatants were transferred to new tubes. Fifiteen µl of each supernatant was saved as input samples and the rest were equally aliquoted for the CPEB1-IP and the IgG-IP (control). The volume of each sample was increased to 300 µl with cLB, 2 µl (2 µg) of the appropriate antibody (anti-CPEB: Abcam, ab73287 and control IgG: Abcam, ab172730) was added to each tube, and samples were incubated on a rotor for 2 hrs at 4°C. Thirty µl washed beads were then added to each tube and samples were incubated on a rotor overnight at 4°C. The beads of each sample were pelleted using a magnetic rack and washed 5 times on a rotor at 4°C with 750 ml wash buffer (WB: 30 mM Tris HCl pH 7.4, 200 mM NaCl, 10 mM MgCl_2_, 0.5% NP-40, 0.25 mM Na_3_VO_4_, 10mM β-glycerophosphate, 1x protease inhibitor, 1 mM DTT, 1M urea, and 1x RNase out) for 10 mins each wash. The beads were then pelleted via a magnetic rack and the WB removed. Two-hundred and fifty µl RLT buffer was added to each sample and the samples were vortexed for 30 secs. RNA extraction was performed following the Rneasy Plus Micro Kit protocol. Samples were eluted in 10 µl of RNase-free water and used for downstream RT-qPCR analysis.

### Real-time quantitative PCR (RT-qPCR)

Extracted RNA was reverse-transcribed using the SuperScript™ III First-Strand Synthesis System with random hexamer primers (Invitrogen, 18080051) and the resulting cDNA was diluted 1:6 with RNase-free water. Gene expression was measured using TaqMan Assays^TM^ and TaqMan^TM^ Fast Advanced Master Mix (ThermoFisher, 4444557). The assays used were: *Astl* (Mm00553165_m1), *Bcl2l10* (Mm00478988_m1), *Ccnb1* (Mm03053893_gH), *Cdk8* (Mm01223097_m1), *Depdc7* (Mm00522683_m1) *Dnmt* (Mm01151063_m1), *Dppa3* (Mm01184198_g1), *Ewsr1* (Mm01191469_g1), *Ing3* (Mm00458324_m1), *Mos* (Mm01700521_g1), *Nlrp5* (Mm01143609_m1), *Obox5* (Mm00773197_gH), *Oosp1* (Mm00504796_m1), *Oosp2* (Mm03015599_m1), *Padi6* (Mm00462201_m1), *Smc4* (Mm00713073_m1), *Tcl1* (Mm00493475_m1), Tiparp (Mm00724822_m1), *Zp1* (Mm00494367_m1), *Zp2* (Mm00442173_m1), and *Zp3* (Mm00442176_m1). Ten µl reactions were run on 384-well plates with the QuantStudio 6 Flex Real-time PCR System. Gene expression was quantified via the 2^-ΔΔCt^ method and statistical analysis was performed via GraphPad Prism 8.

### Construction of florescent protein reporters

The 3’ UTR sequences of *Ccnb1*, *Ccnb2 short, Ewsr1*, *Mos*, *Oosp1*, *Oosp2*, *Smc4*, and *Zp2* were retrieved from the RNA-Seq .bam files using the Integrative Genomics Viewer (Supplementary Table 1). Primers were used to amplify the target 3’ UTRs from oocyte cDNA and portions of YFP-containing vector. Using the Choo-Choo Cloning^TM^ Kit (MCLAB, CCK-20), the PCR fragments were fused together and transfected into competent 5-α *E. coli* cells. The DNA plasmids of ampicillin-resistant bacteria were extracted using the QIAprep Spin Miniprep Kit (Qiagen, 27106) and the sequences confirmed via DNA sequencing. The extracted plasmid was then linearized using a forward primer upstream of the YFP sequence and a reverse primer to the 3’UTR with 20 additional thymine residues. The PCR product was then transcribed *in vitro* using the mMESSAGE mMACHINE T7 Transcription Kit (Invitrogen, AM1344) and the resulting cRNA was purified using the MEGAclear™ Transcription Clean-Up Kit (Invitrogen, AM1908); cRNA were eluted in RNase-free water and kept at −80°C. The mCherry reporter was similarly produced. However, the message contained no 3’UTR, but instead was polyadenylated (150-200 nts) using the Poly(A) Tailing Kit (Invitrogen, AM1350).

### Oocyte microinjection

Oocytes were collected as described and allowed to recover in αC media for two hrs, after which they were transferred into HC media for microinjection. Oocytes were injected with 5-10 pl of a 12.5 ng/µl solution of the YFP reporter of interest mixed with *mCherry* mRNA and allowed to recover in αC media for the specified amount of time before live cell imaging.

### Live cell imaging and fluorescence microscopy

Live cell imaging experiments were performed using a Nikon Eclipse T2000-E equipped with mobile stage and environmental chamber at 37°C and 5% CO_2_. Filter set: dichroic mirror YFP/CFP/mCherry 69008BS; Ypet channel (Ex: S500/20x 49057; Em: D535/30m 47281), mCherry channel (Ex: 580/25x 49829; Em: 632/60m). Images were processed and fluorescence was quantified using MetaMorph.

### Poly(A) tail length (PAT) assay

This assay was performed as previously described(Yang et al., 2017).

### Histology

Ovaries were dissected from eight week-old female mice, fixed in Bouin’s solution, and preserved in 70% ethanol. Tissues were then processed, cut at 8 µm, and stained (H&E) by the Cancer Center Tissue Core at UCSF.

### Western blot

Oocytes were collected in 0.1% PVP in DPBS and boiled for 3 mins at 95°C in 1x Laemmli Sample Buffer (Bio-Rad, 161-0747) supplemented with β-mercaptoethanol, proteinase inhibitor, and phosphatase inhibitor. Samples were resolved on a 10% Laemmli gel and semi-dry transferred onto supported nitrocellulose membranes, 0.2 µm (Bio-Rad, 1620097). Membranes were incubated in 5% blocking buffer for 1 hr then incubated for 18 hrs in primary antibody at 4°C. The antibodies used: 1:1000 rabbit anti-CPEB1 (Abcam, ab73287). The membrane was then washed in 1x TBST, incubated in the appropriate secondary antibodies, 1:20,000 rabbit IgG (GE Healthcare, NA934V) for 2 hrs, and washed again in 1x TBST. Clarity Western ECL substrate (Bio-Rad, 1705061) was then used to develop the membrane. All washes were done four times each round in TBST for 10 mins each wash.

### Materials availability

Further information and requests for resources and reagents should be directed to and will be fulfilled by the Lead Contact, Marco Conti.

### Data and code availability

Raw sequences and TMM-normalized CPM values from the RiboTag/RNA-Seq experiment will be available on UCSF Box. Scripts used for statistical analysis of the RiboTag/RNA-Seq data and those used to calculate CPE and PAS distances will be available on UCSF Box.

## References

Bailey, T.L., Boden, M., Buske, F.A., Frith, M., Grant, C.E., Clementi, L., Ren, J.Y., Li, W.W., and Noble, W.S. (2009). MEME SUITE: tools for motif discovery and searching. Nucleic Acids Research 37, W202–W208.

Ballantyne, S., Daniel, D.L., and Wickens, M. (1997). A dependent pathway of cytoplasmic polyadenylation reactions linked to cell cycle control by c-mos and CDK1 activation. Molecular Biology of the Cell 8, 1633–1648.

Barnard, D.C., Ryan, K., Manley, J.L., and Richter, J.D. (2004). Symplekin and xGLD-2 are required for CPEB-mediated cytoplasmic polyadenylation. Cell 119, 641–651.

Beaudoing, E., Freier, S., Wyatt, J.R., Claverie, J.M., and Gautheret, D. (2000). Patterns of variant polyadenylation signal usage in human genes. Genome Research 10, 1001–1010.

Bolger, A.M., Lohse, M., and Usadel, B. (2014). Trimmomatic: a flexible trimmer for Illumina sequence data. Bioinformatics 30, 2114–2120.

Calderone, V., Gallego, J., Fernandez-Miranda, G., Garcia-Pras, E., Maillo, C., Berzigotti, A., Mejias, M., Bava, F.A., Angulo-Urarte, A., Graupera, M., et al. (2016). Sequential Functions of CPEB1 and CPEB4 Regulate Pathologic Expression of Vascular Endothelial Growth Factor and Angiogenesis in Chronic Liver Disease. Gastroenterology 150, 982-+.

Chen, J., Melton, C., Suh, N., Oh, J.S., Horner, K., Xie, F., Sette, C., Blelloch, R., and Conti, M. (2011). Genome-wide analysis of translation reveals a critical role for deleted in azoospermia-like (Dazl) at the oocyte-to-zygote transition. Genes & Development 25, 755–766.

Chen, T.P., and Dent, S.Y.R. (2014). Chromatin modifiers and remodellers: regulators of cellular differentiation. Nature Reviews Genetics 15, 93–106.

Cheng, J., Maier, K.C., Avsec, Z., Rus, P., and Gagneur, J. (2017). Cis-regulatory elements explain most of the mRNA stability variation across genes in yeast. Rna 23, 1648–1659.

Chiu, T.P., Yang, L., Zhou, T.Y., Main, B.J., Parker, S.C.J., Nuzhdin, S.V., Tullius, T.D., and Rohs, R. (2015). GBshape: a genome browser database for DNA shape annotations. Nucleic Acids Research 43, D103–D109.

Clarke, H.J. (2012). Post-transcriptional control of gene expression during mouse oogenesis. Results and problems in cell differentiation 55, 1–21.

Conti, M., and Franciosi, F. (2018). Acquisition of oocyte competence to develop as an embryo: integrated nuclear and cytoplasmic events. Human Reproduction Update 24, 245–266.

Evans, T., Rosenthal, E.T., Youngblom, J., Distel, D., and Hunt, T. (1983). CYCLIN - A PROTEIN SPECIFIED BY MATERNAL MESSENGER-RNA IN SEA-URCHIN EGGS THAT IS DESTROYED AT EACH CLEAVAGE DIVISION. Cell 33, 389–396.

Grant, C.E., Bailey, T.L., and Noble, W.S. (2011). FIMO: scanning for occurrences of a given motif. Bioinformatics 27, 1017–1018.

Han, S.J., Martins, J.P.S., Yang, Y., Kang, M.K., Daldello, E.M., and Conti, M. (2017). The Translation of Cyclin B1 and B2 is Differentially Regulated during Mouse Oocyte Reentry into the Meiotic Cell Cycle. Scientific Reports 7.

Horvat, F., Fulka, H., Jankele, R., Malik, R., Jun, M., Solcova, K., Sedlacek, R., Vlahovicek, K., Schultz, R.M., and Svoboda, P. (2018). Role of Cnot6l in maternal mRNA turnover. Life Science Alliance 1.

Huang, D.W., Sherman, B.T., and Lempicki, R.A. (2009a). Bioinformatics enrichment tools: paths toward the comprehensive functional analysis of large gene lists. Nucleic Acids Research 37, 1–13.

Huang, D.W., Sherman, B.T., and Lempicki, R.A. (2009b). Systematic and integrative analysis of large gene lists using DAVID bioinformatics resources. Nature Protocols 4, 44–57.

Ivanova, I., Much, C., Di Giacomo, M., Azzi, C., Morgan, M., Moreira, P.N., Monahan, J., Carrieri, C., Enright, A.J., and O’Carroll, D. (2017). The RNA m(6)A Reader YTHDF2 Is Essential for the Post-transcriptional Regulation of the Maternal Transcriptome and Oocyte Competence. Molecular Cell 67, 1059-+.

Jacobson, A., and Peltz, S.W. (1996). Interrelationships of the pathways of mRNA decay and translation in eukaryotic cells. Annual Review of Biochemistry 65, 693–739.

Jonas, S., and Izaurralde, E. (2015). NON-CODING RNA Towards a molecular understanding of microRNA-mediated gene silencing. Nature Reviews Genetics 16, 421–433.

Kim, D., Landmead, B., and Salzberg, S.L. (2015). HISAT: a fast spliced aligner with low memory requirements. Nature Methods 12, 357–U121.

Kimble, J., and Crittenden, S.L. (2007). Controls of germline stem cells, entry into meiosis, and the Sperm/Oocyte decision in Caenorhabditis elegans. Annual Review of Cell and Developmental Biology 23, 405–433.

Klemm, S.L., Shipony, Z., and Greenleaf, W.J. (2019). Chromatin accessibility and the regulatory epigenome. Nature Reviews Genetics 20, 207–220.

Li, H., Handsaker, B., Wysoker, A., Fennell, T., Ruan, J., Homer, N., Marth, G., Abecasis, G., Durbin, R., and Genome Project Data, P. (2009). The Sequence Alignment/Map format and SAMtools. Bioinformatics 25, 2078–2079.

Liu, Y.S., Lu, X.K., Shi, J.C., Yu, X.J., Zhang, X.X., Zhu, K., Yi, Z.H., Duan, E.K., and Li, L. (2016). BTG4 is a key regulator for maternal mRNA clearance during mouse early embryogenesis. Journal of Molecular Cell Biology 8, 366–368.

Martins, J.P.S., Liu, X., Oke, A., Arora, R., Franciosi, F., Viville, S., Laird, D.J., Fung, J.C., and Conti, M. (2016). DAZL and CPEB1 regulate mRNA translation synergistically during oocyte maturation. Journal of Cell Science 129, 1271–1282.

Mendez, R., Barnard, D., and Richter, J.D. (2002). Differential rnRNA translation and rneiotic progression require Cdc2-mediated CPEB destruction. Embo Journal 21, 1833–1844.

Mendez, R., Hake, L.E., Andresson, T., Littlepage, L.E., Ruderman, J.V., and Richter, J.D. (2000a). Phosphorylation of CPE binding factor by Eg2 regulates translation of c-mos mRNA. Nature 404, 302–307.

Mendez, R., Murthy, K.G.K., Ryan, K., Manley, J.L., and Richter, J.D. (2000b). Phosphorylation of CPEB by Eg2 mediates the recruitment of CPSF into an active cytoplasmic polyadenylation complex. Molecular Cell 6, 1253–1259.

Mendez, R., and Richter, J.D. (2001). Translational control by CPEB: A means to the end. Nature Reviews Molecular Cell Biology 2, 521–529.

Morgan, M., Much, C., DiGiacomo, M., Azzi, C., Ivanova, I., Vitsios, D.M., Pistolic, J., Collier, P., Moreira, P.N., Benes, V., et al. (2017). mRNA 3’ uridylation and poly(A) tail length sculpt the mammalian maternal transcriptome. Nature 548, 347-+.

Paillisson, A., Dade, S., Callebaut, I., Bontoux, M., Dalbies-Tran, R., Vaiman, D., and Monget, P. (2005). Identification, characterization and metagenome analysis of oocyte-specific genes organized in clusters in the mouse genome. Bmc Genomics 6.

Park, J.E., Yi, H., Kim, Y., Chang, H., and Kim, V.N. (2016). Regulation of Poly(A) Tail and Translation during the Somatic Cell Cycle. Molecular Cell 62, 462–471.

Pasternak, M., Pfender, S., Santhanam, B., and Schuh, M. (2016). The BTG4 and CAF1 complex prevents the spontaneous activation of eggs by deadenylating maternal mRNAs. Open Biology 6.

Pique, M., Lopez, J.M., Foissac, S., Guigo, R., and Mendez, R. (2008). A combinatorial code for CPE-mediated translational control. Cell 132, 434–448.

Racki, W.J., and Richter, J.D. (2006). CPEB controls oocyte growth and follicle development in the mouse. Development 133, 4527–4537.

Radford, H.E., Meijer, H.A., and de Moor, C.H. (2008). Translational control by cytoplasmic polyadenylation in Xenopus oocytes. Biochimica Et Biophysica Acta-Gene Regulatory Mechanisms 1779, 217–229.

Richter, J.D. (2007). CPEB: a life in translation. Trends in Biochemical Sciences 32, 279–285.

Richter, J.D., and Lasko, P. (2011). Translational Control in Oocyte Development. Cold Spring Harbor Perspectives in Biology 3.

Rissland, O.S. (2017). The organization and regulation of mRNA-protein complexes. Wiley Interdisciplinary Reviews-Rna 8.

Rissland, O.S., Subtelny, A.O., Wang, M., Lugowski, A., Nicholson, B., Laver, J.D., Sidhu, S.S., Smibert, C.A., Lipshitz, H.D., and Bartel, D.P. (2017). The influence of microRNAs and poly(A) tail length on endogenous mRNA-protein complexes. Genome Biology 18.

Ritchie, M.E., Phipson, B., Wu, D., Hu, Y.F., Law, C.W., Shi, W., and Smyth, G.K. (2015). limma powers differential expression analyses for RNA-sequencing and microarray studies. Nucleic Acids Research 43.

Robinson, M.D., McCarthy, D.J., and Smyth, G.K. (2010). edgeR: a Bioconductor package for differential expression analysis of digital gene expression data. Bioinformatics 26, 139–140.

Rueden, C.T., Schindelin, J., Hiner, M.C., DeZonia, B.E., Walter, A.E., Arena, E.T., and Eliceiri, K.W. (2017). ImageJ2: ImageJ for the next generation of scientific image data. Bmc Bioinformatics 18.

Seydoux, G., and Braun, R.E. (2006). Pathway to totipotency: Lessons from germ cells. Cell 127, 891–904.

Sheets, M.D., Fox, C.A., Hunt, T., Vandewoude, G., and Wickens, M. (1994). THE 3’-UNTRANSLATED REGIONS OF C-MOS AND CYCLIN MESSENGER-RNAS STIMULATE TRANSLATION BY REGULATING CYTOPLASMIC POLYADENYLATION. Genes & Development 8, 926–938.

Simon, R., and Richter, J.D. (1994). FURTHER ANALYSIS OF CYTOPLASMIC POLYADENYLATION IN XENOPUS EMBRYOS AND IDENTIFICATION OF EMBRYONIC CYTOPLASMIC POLYADENYLATION ELEMENT-BINDING PROTEINS. Molecular and Cellular Biology 14, 7867–7875.

Su, Y.Q., Sugiura, K., Woo, Y., Wigglesworth, K., Kamdar, S., Affourtit, J., and Eppig, J.J. (2007). Selective degradation of transcripts during meiotic maturation of mouse oocytes. Developmental Biology 302, 104–117.

Subtelny, A.O., Eichhorn, S.W., Chen, G.R., Sive, H., and Bartel, D.P. (2014). Poly(A)-tail profiling reveals an embryonic switch in translational control. Nature 508, 66-+.

Tadros, W., and Lipshitz, H.D. (2005). Setting the stage for development: mRNA translation and stability during oocyte maturation and egg activation in Drosophila. Developmental Dynamics 232, 593–608.

Tay, J., Hodgman, R., and Richter, J.D. (2000). The control of cyclin B1 mRNA translation during mouse oocyte maturation. Developmental Biology 221, 1–9.

Tay, J., and Richter, J.D. (2001). Germ cell differentiation and synaptonemal complex formation are disrupted in CPEB knockout mice. Developmental Cell 1, 201–213.

Wang, S.F., Kou, Z.H., Jing, Z.Y., Zhang, Y., Guo, X.Z., Dong, M.Q., Wilmut, I., and Gao, S.R. (2010). Proteome of mouse oocytes at different developmental stages. Proceedings of the National Academy of Sciences of the United States of America 107, 17639–17644.

Wang, X., Lu, Z.K., Gomez, A., Hon, G.C., Yue, Y.N., Han, D.L., Fu, Y., Parisien, M., Dai, Q., Jia, G.F., et al. (2014). N-6-methyladenosine-dependent regulation of messenger RNA stability. Nature 505, 117-+.

Wickham, H. (2009). ggplot2 Elegant Graphics for Data Analysis Introduction. Ggplot2: Elegant Graphics for Data Analysis, 1-+.

Wu, X.Y., and Brewer, G. (2012). The regulation of mRNA stability in mammalian cells: 2.0. Gene 500, 10–21.

Yang, Y., Yang, C.-R., Han, S.J., Daldello, E.M., Cho, A., Martins, J.P.S., Xia, G., and Conti, M. (2017). Maternal mRNAs with distinct 3’ UTRs define the temporal pattern of Ccnb1 synthesis during mouse oocyte meiotic maturation. Genes & Development 31, 1302–1307.

Yartseva, V., and Giraldez, A.J. (2015). The Maternal-to-Zygotic Transition During Vertebrate Development: A Model for Reprogramming. Maternal-to-Zygotic Transition 113, 191–232.

Yu, C., Ji, S.Y., Sha, Q.Q., Dang, Y.J., Zhou, J.J., Zhang, Y.L., Liu, Y., Wang, Z.W., Hu, B.Q., Sun, Q.Y., et al. (2016). BTG4 is a meiotic cell cycle-coupled maternal-zygotic transition licensing factor in oocytes. Nature Structural & Molecular Biology 23, 387–394.

