## Supplemental material for "Genome-wide analysis reveals a switch in the translational program upon oocyte meiotic resumption"

### Supplemental Figure 1

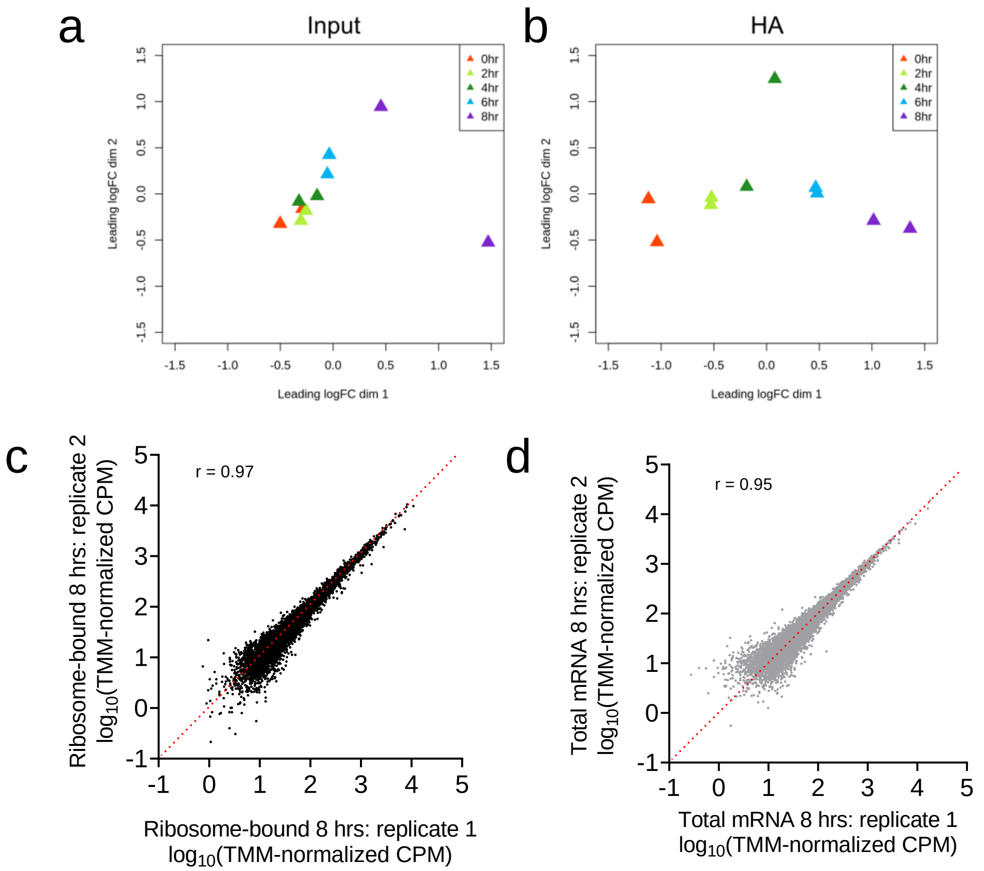


e

| HA-8 hrs | CONSTITUTIVE | DOWN | UP |
|---|---|---|---|
| FDR ≤0.05 | 4284 | 1722 | 1537 |
| FDR ≤0.01 | 5104 | 1263 | 1176 |
| FDR ≤0.001 | 5871 | 815 | 857 |
| **FDR ≤0.05, -1 ≥ LFC ≥1** | **5580** | **1092** | **871** |
| FDR ≤0.01, -1 ≥ LFC ≥1 | 5858 | 913 | 772 |
| FDR ≤0.001, -1 ≥ LFC ≥1 | 6236 | 658 | 649 |

f

| Input-8 hrs | NS | Decrease | Increase |
|---|---|---|---|
| FDR ≤0.05 | 7473 | 322 | 122 |
| FDR ≤0.01 | 7676 | 194 | 47 |
| FDR ≤0.001 | 7782 | 119 | 16 |
| **FDR ≤0.05, -1 ≥ LFC ≥1** | **7666** | **218** | **33** |
| FDR ≤0.01, -1 ≥ LFC ≥1 | 7739 | 162 | 16 |
| FDR ≤0.001, -1 ≥ LFC ≥1 | 7799 | 110 | 8 |

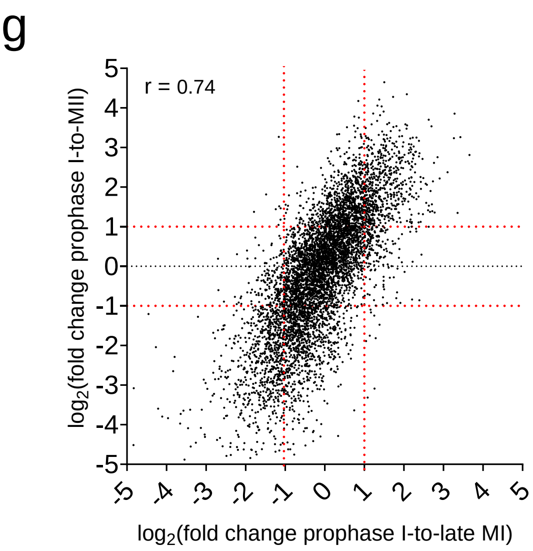

**Supp. Fig. 1. Validation of RiboTag/RNA-Seq method and summary of data**

**a-b)** Multidimensional scaling plots for input (a) and HA-IP (b) samples. TMM-normalized CPM data and the plotMDS function from the limma package were used to generate the plots. Input data cluster together well except for the 8 hrs samples, while the HA-immunoprecipitated groups display clustering according to the sampling time. This suggests changes in total mRNA levels at late MI as well as time-dependent changes in ribosome association of maternal messages. **c-d)** Correlation between replicate late MI samples. TMM-normalized CPM values of the duplicate total mRNA samples at late MI are highly correlated with one another (c; Pearson correlation: r = 0.97); similarly, high correlation is present for the duplicate ribosome-bound mRNA samples (d; Pearson correlation: r = 0.95). **e-f)** Differential gene expression analysis. Pair-wise statistical analysis was performed on TMM-normalized CPM values comparing late MI to prophase I. Listed are the number of maternal mRNAs differentially expressed throughout the time course experiment for ribosome-bound (e) and total mRNA (f) samples using different fold change (FC) and false discovery rate (FDR) thresholds; NS: not significant. **g)** Correlation between changes in ribosome loading at late MI and MII. Changes in translation between prophase I and MII obtained from published polysome array data[27, 42] were correlated with changes in translation between prophase I and late MI obtained from the RiboTag/RNA-Seq dataset. There is a strong, positive correlation between translatome changes in late MI and translatome changes in MII (Pearson correlation: r = 0.74), confirming consistent patterns of differential translation throughout meiotic maturation.

### Supplemental Figure 2

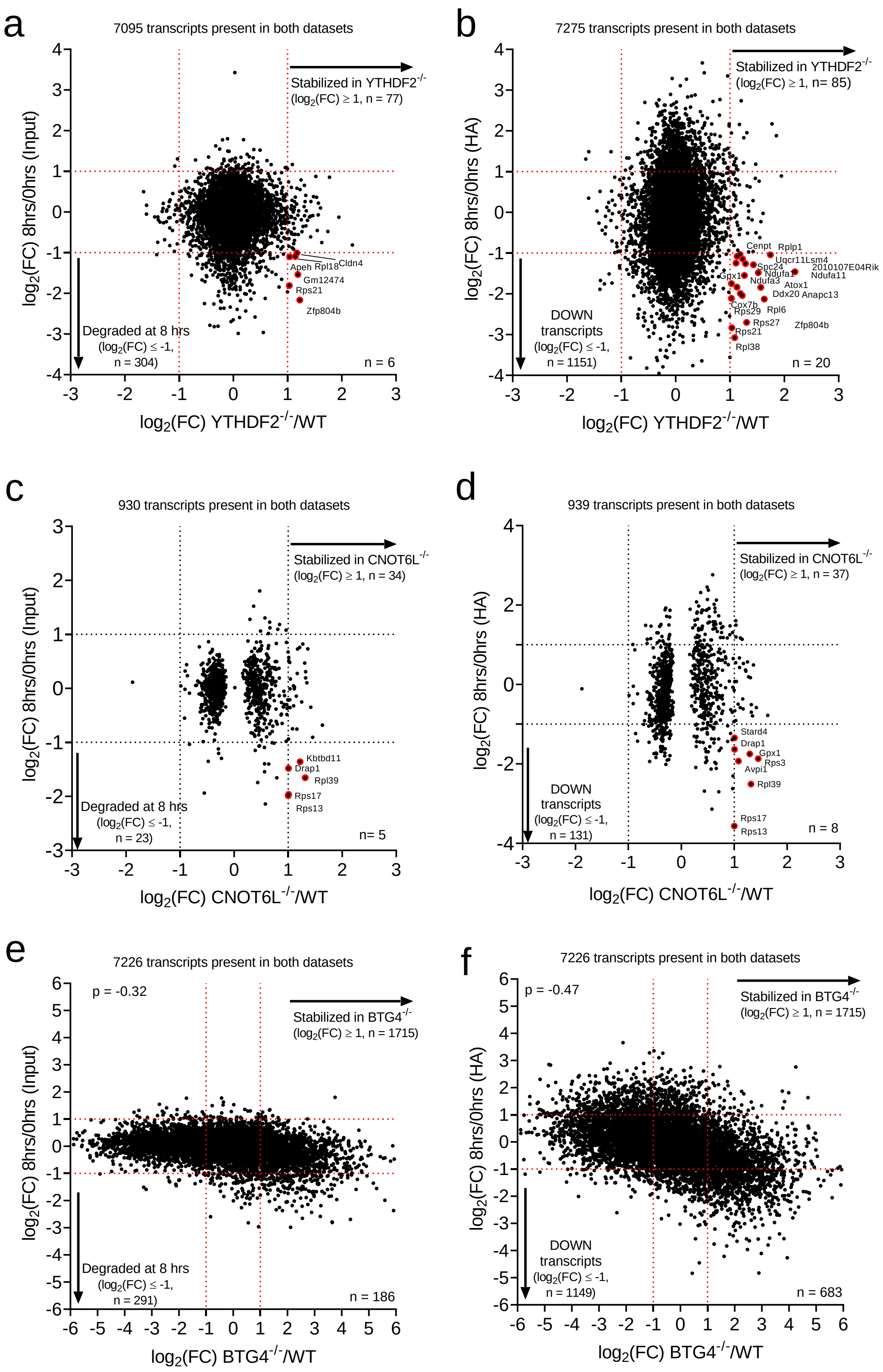

**Supp. Fig. 2. Comparison of transcripts undergoing degradation or translation repression during meiosis with those stabilized in YTHDF2$^{-/-}$, CNOT6L$^{-/-}$, and BTG4$^{-/-}$ oocytes.** The mRNAs present in our RNA-Seq dataset was compared pairwise to YTHDF2$^{-/-}$[22] (a-b), CNOT6L$^{-/-}$[23] (c-d), and BTG4$^{-/-}$[25] (e-f) datasets. The number of genes common to both datasets is included in the figure. Fold changes in total mRNA levels between 0 and 8 hrs were compared with mRNAs that are stabilized in YTHDF2$^{-/-}$ (a), CNOT6L$^{-/-}$ (c), and BTG4$^{-/-}$ (e) oocytes. Similarly, fold changes in the ribosome-bound mRNA levels between 0 and 8 hrs were compared to mRNAs that are stabilized in YTHDF2$^{-/-}$ (b), CNOT6L$^{-/-}$ (d), and BTG4$^{-/-}$ (f) oocytes. There is a significant overlap between mRNAs that are translationally repressed (n = 683 of 1149, 59%) or destabilized by late MI (n = 186 of 291, 64%) with those degraded by MII in a BTG4-dependent manner.

### Supplemental Figure 3

#### *Bub1b*

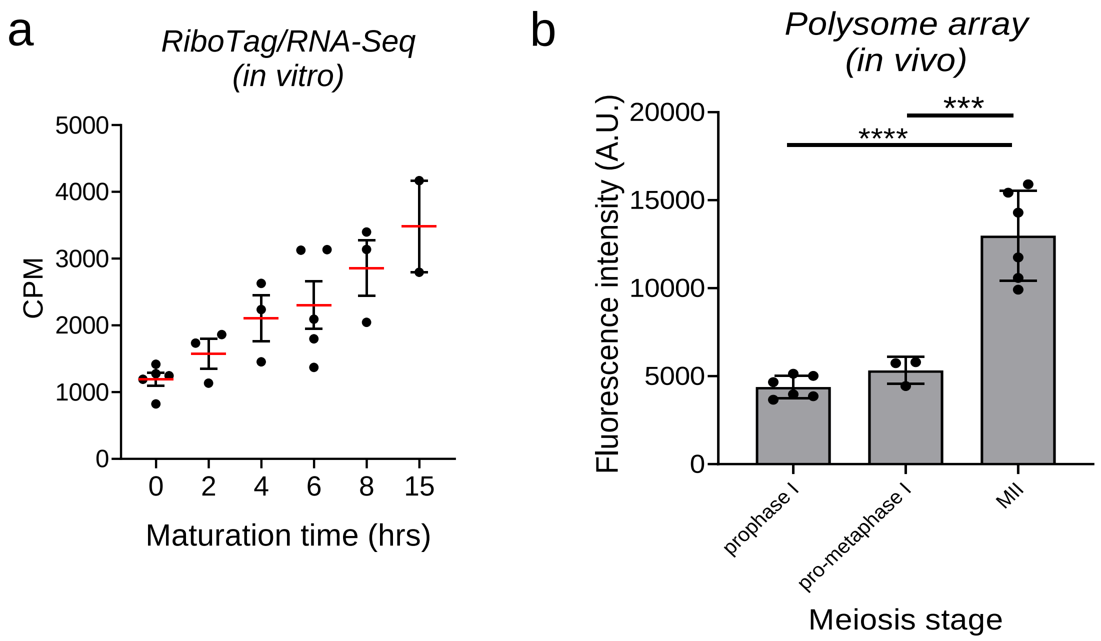


#### *Cdk1*

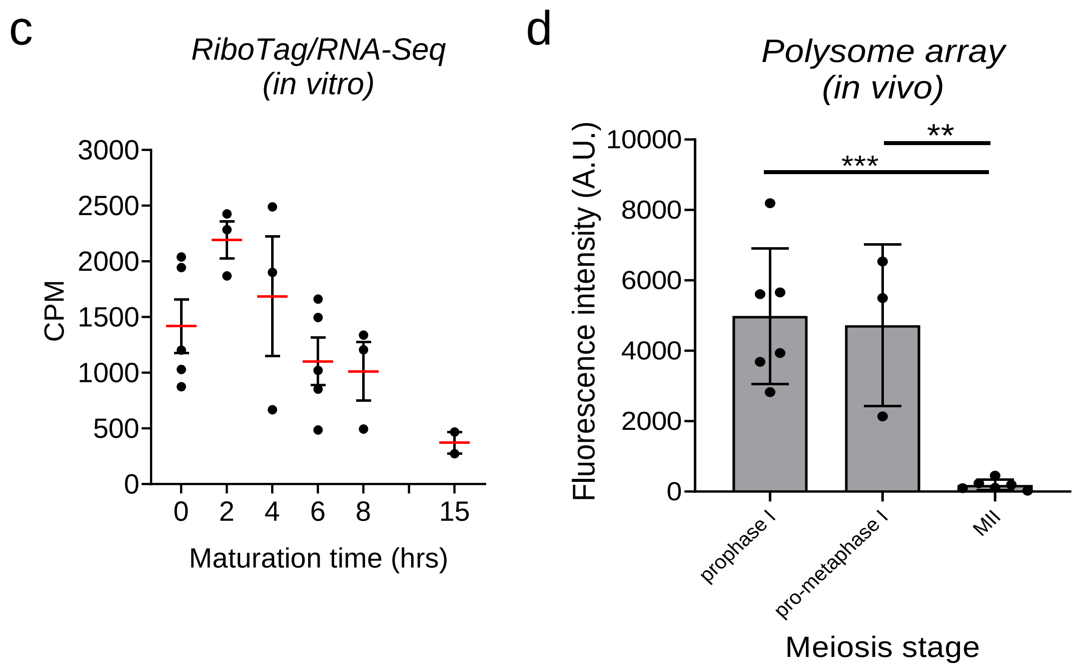

**Supp. Fig. 3. Changes in mRNA translation are consistent across multiple biological replicates**

TMM-normalized CPM values from the two replicates of our RiboTag/RNA-Seq experiment were compared to a single point of a previous pilot RiboTag/RNA-Seq study[21], a 0 hrs/6 hrs dataset that is under review, a published GV/MI/MII dataset[42], and a published GV/MII dataset[27]. No cross study normalizations nor corrections for batch effects were applied. Translation of the SAC checkpoint kinase *Bub1b* increased during meiotic resumption (a-b), while translation of *Cdk1* consistently decreased (c-d).

### Supplemental Figure 4

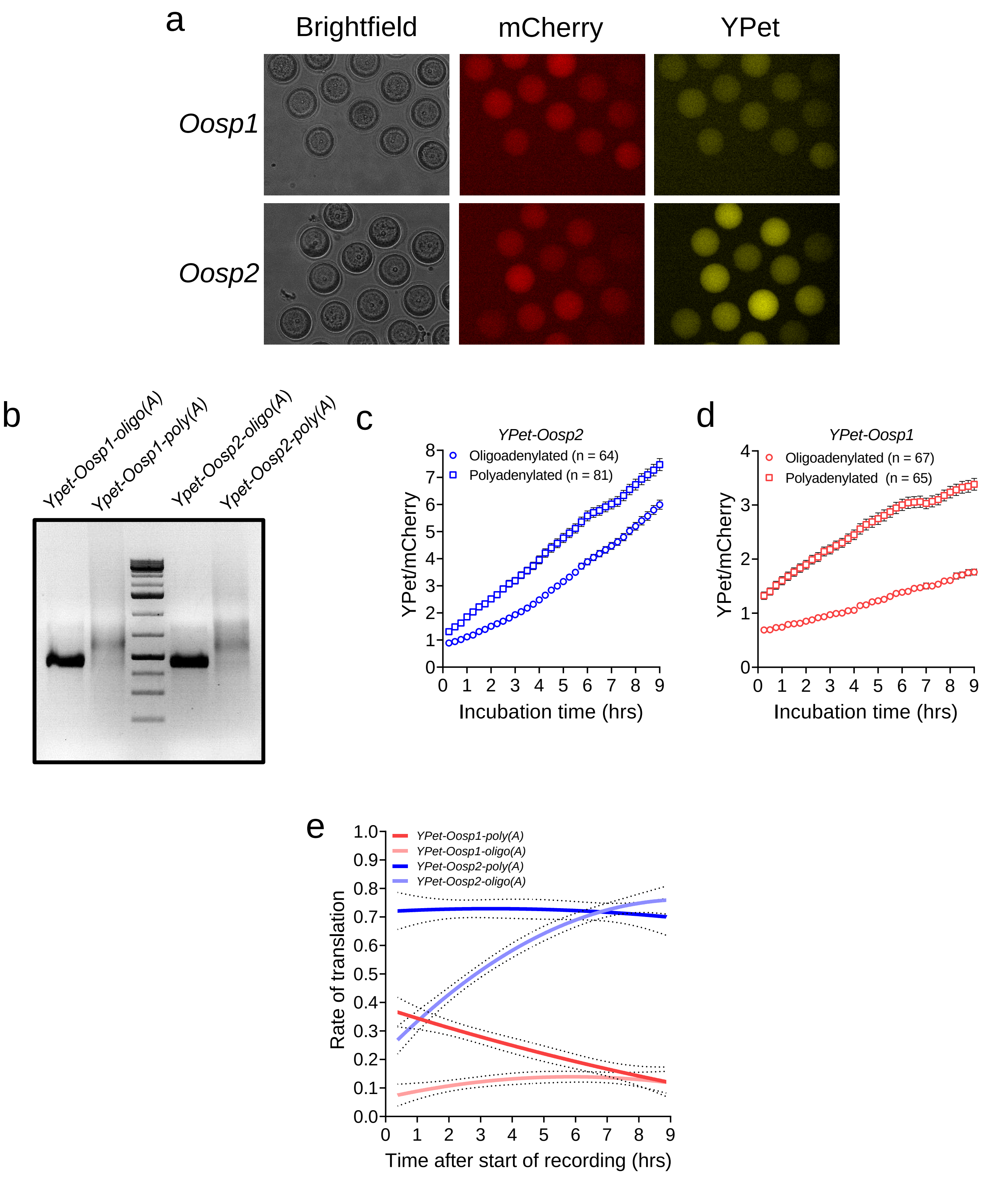

**Supp. Fig. 4. Translation of the *Oosp1* and *Oosp2* oligoadenylated or polyadenylated reporters**

**a)** Example images of oocytes under brightfield, mCherry, or YPet wavelengths. Images were taken 10 hrs after the start of recording and are from experiments reported in Fig. 5e. Oocytes were injected with *Oosp1* and *Oosp2* YPet reporters and *mCherry*. **b)** Gel electrophoresis images of *in vitro* transcribed *YPet-Oosp1* and *YPet-Oosp2* mRNAs. Reporters were either oligoadenylated or polyadenylated. Aliquots of each message were loaded onto a 2% agarose gel with ethidium bromide and developed under UV light. Polyadenylated messages displayed heavier bands with increased smearing. **c)** YPet accumulation in GV-arrested oocytes injected with either oligoadenylated or polyadenylated *YPet-Oosp2* reporter. The data are reported as YPet signal corrected by averaged mCherry signal. Each point is the mean ± SEM of individual oocyte traces and the total number of oocytes analyzed is in parentheses. **d)** YPet accumulation in GV-arrested oocytes injected with either oligoadenylated or polyadenylated *YPet-Oosp1* reporter. The data are reported as YPet signal corrected by averaged mCherry signal. Each point is the mean ± SEM of individual oocyte traces and the total number of oocytes analyzed is in parentheses. **e)** Rates of reporter translation over time in GV-arrested oocytes. Rates of translation were calculated as the first derivative of the YPet/mCherry traces (Supp. Fig. 2c and d) and the resulting curves were fitted to a quadratic curve. The 95% confidence intervals (dashed lines) are plotted along with the fitted curves (solid lines). In GV-arrested oocytes, the rates of translation of *YPet-Oosp2-poly(A)* (blue) remains steady during the 9 hr incubation, while translation of *YPet-Oosp2-oligo(A)* (light blue) increases to levels similar to those of its polyadenylated counterpart. Translation of *YPet-Oosp1-oligo(A)* (light red), however, remains steady during this time and translation of *YPet-Oosp1-poly(A)* decreases to levels similar to those of its oligoadenylated counterpart (red).

### Supplemental Figure 5

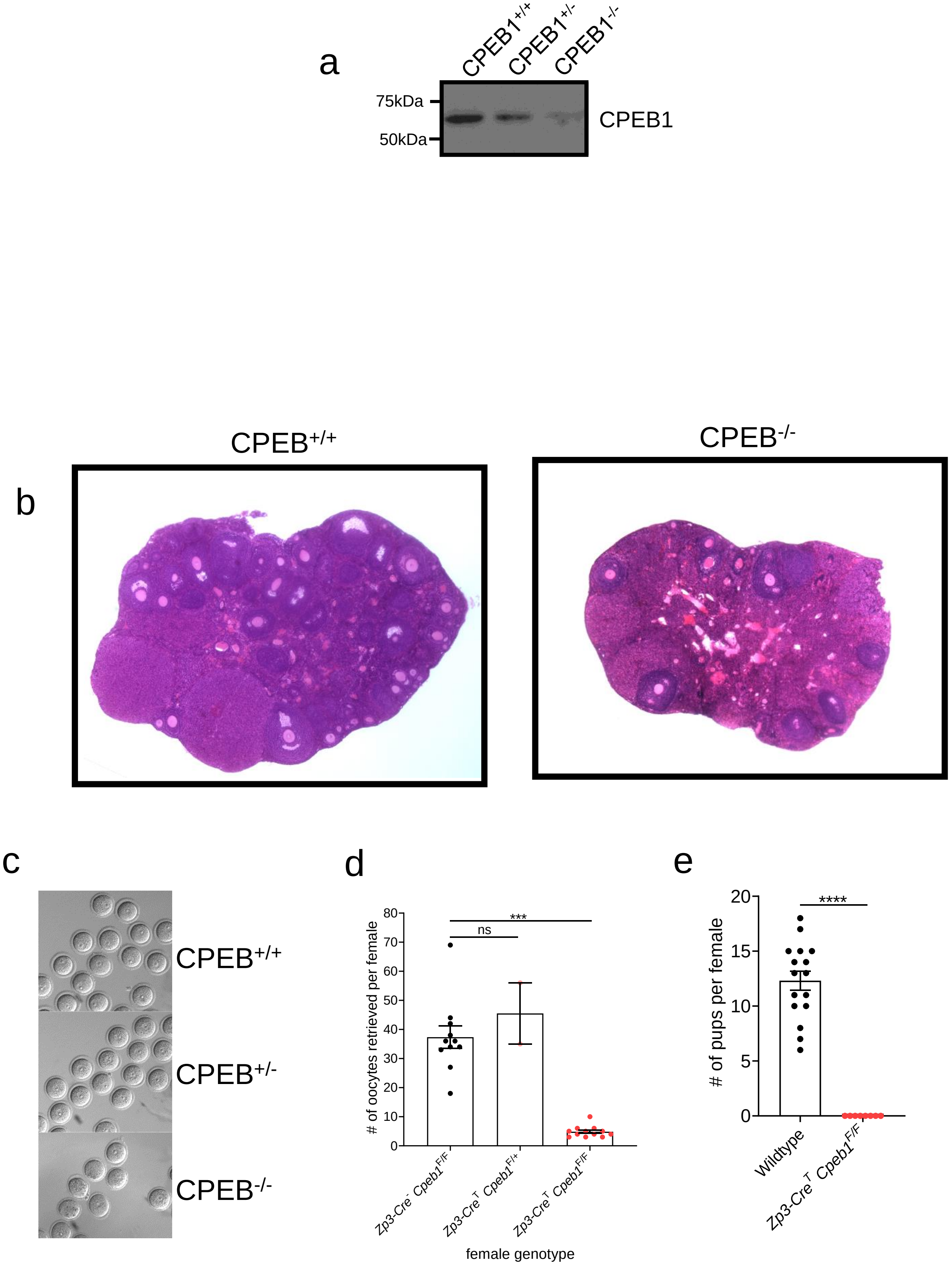

**Supp. Fig. 5. Phenotypes of oocyte-specific CPEB1$^{-/-}$ female mice**

We used a *Zp3* driven Cre recombinase strategy to delete *Cpeb1* specifically in the oocytes. **a)** Image of Western blot membrane of mouse oocytes using CPEB1-antibodies. Thirty CPEB1$^{+/+}$, CPEB1$^{+/-}$, and CPEB1$^{-/-}$ GV-arrested oocytes were used for this experiment. A dose-dependent depletion of CPEB1 in the oocytes is observed according to the genotype. **b)** Histological sections of wild type and *Zp3-Cre*$^{T}$ *Cpeb1*$^{F/F}$ mouse ovaries. Images of representative ovaries from eight-week-old wild type and transgenic mice are reported. *Zp3-Cre*$^{T}$ *Cpeb1*$^{F/F}$ ovaries contain antral follicles although with decreased numbers when compared to wild type ovaries. **c)** Brightfield images of CPEB1$^{+/+}$, CPEB1$^{+/-}$, and CPEB1$^{-/-}$ oocytes. Ovaries were dissected in media containing cilostamide. While all CPEB$^{+/+}$ and CPEB$^{+/-}$ oocytes retrieved were arrested in prophase I and displayed a visible germinal vesicle, some CPEB$^{-/-}$ oocytes collected had already undergone GVBD, suggesting spontaneous maturation. **d)** Number of oocytes collected per female. Wild type, *Zp3-Cre*$^{T}$ *Cpeb1*$^{F/+}$, and *Zp3-Cre*$^{T}$ *Cpeb1*$^{F/F}$ were primed with PMGS and ovaries were dissected out 44 hrs later. Mean ± SEM is reported. Significantly fewer oocytes were collected from individual CPEB1 knockout female mice (mean = 4, n = 11) as compared to wild type (mean = 37, n = 2) or heterozygous females (mean = 45, n = 12). Statistical significance was evaluated by Kruskal-Wallis test; ***: $p = 0.0001$; ns: not significant **e)** Number of pups from CPEB1 KO mating. Wild type and *Zp3-Cre*$^{T}$ *Cpeb1*$^{F/F}$ two months of age were mated with wild type males of comparable age and monitored for four months. Mean ± SEM is reported. *Zp3-Cre*$^{T}$ *Cpeb1*$^{F/F}$ females were completely infertile (mean = 0, n = 8), while wild type females successfully produced litters (mean = 12, n = 16). Statistical significance was evaluated by Mann-Whitney test; ****: $p < 0.0001$. Previous reports of mid-gestation global CPEB knockout female mice have ovaries that contain oocytes arrested at the pachytene stage and become devoid of oocytes in adult mice[59]. In conditional knockdown females, when *Cpeb*-targeted siRNA is expressed in growing dictyate stage oocytes via the *Zp3* promoter, gametes are observed in the ovaries of six week-old females[60]. These knockdown oocytes displayed several developmental phenotypes including premature maturation, improper chromosome alignment at the metaphase plate, and abnormal polar bodies, resulting in significantly reduced fertility in the transgenic females. Our data with the conditional KO are comparable to these previous observations.

### Supplemental Figure 6

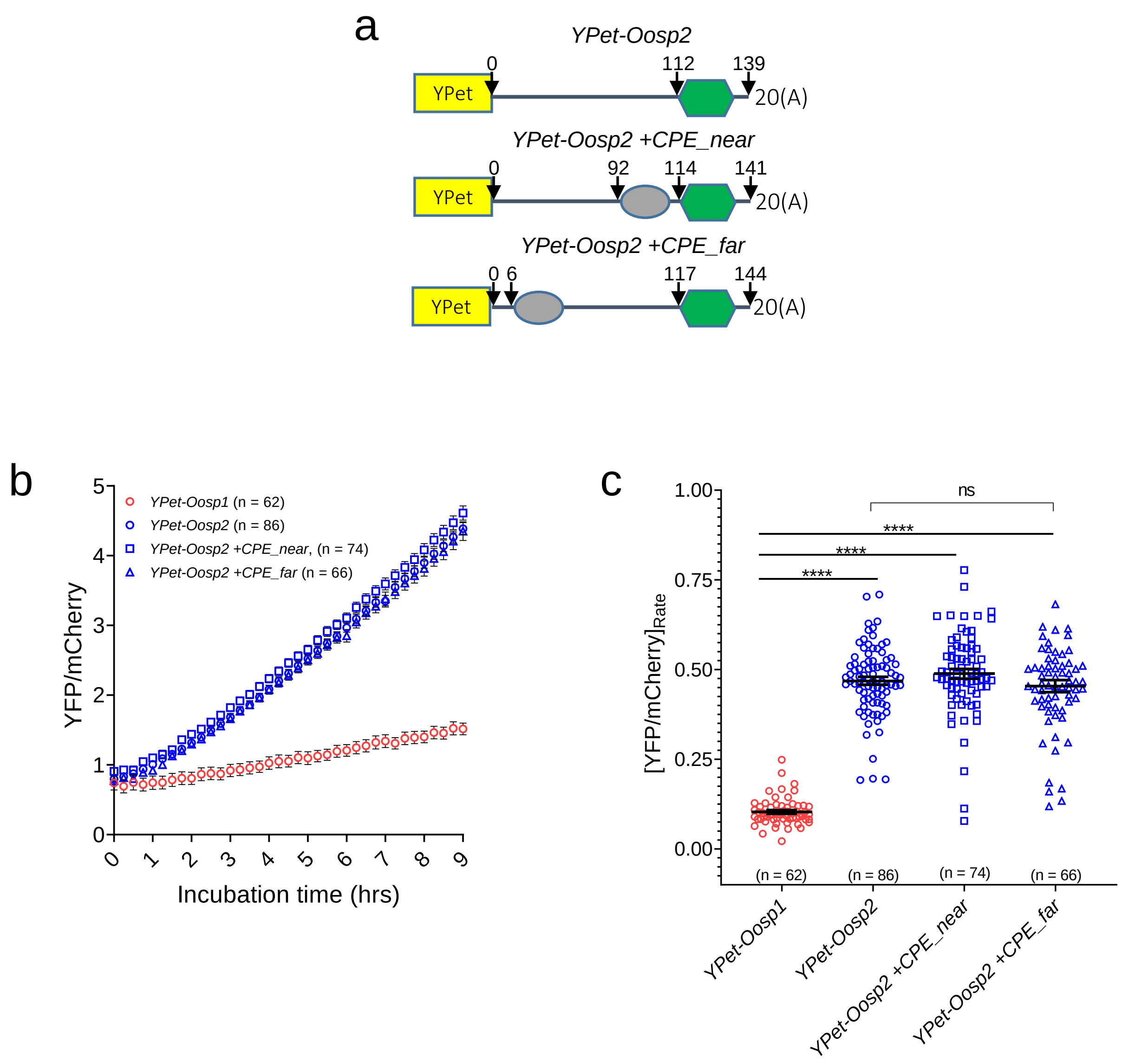

**Supp. Fig. 6. Inclusion of CPEs to the 3'UTR is not sufficient to repress *Oosp2* translation in GV-arrested oocytes.**

**a)** YPet reporters for various *Oosp2* 3'UTRs. The wild type 3'UTR of *Oosp2* was fused downstream of a YPet protein ORF (yellow rectangle) and the message was oligoadenylated (*YPet-Oosp2*). Mutant 3'UTRs were obtained by adding a single CPE (TTTTAT) either near or far from the PAS (*YPet-Oosp2 +CPE_near* and *YPet-Oosp2 +CPE_near*, respectively). Relevant nucleotide positions relative to the start of the 3'UTR are displayed. **b)** Accumulation of wild type *Oosp1*, wild type *Oosp2*, and mutant *Oosp2* reporters in GV-arrested oocytes. GV-arrested oocytes were collected and microinjected with the reporters described in Supplementary Fig. 6a along with polyadenylated *mCherry* mRNA. Oocytes were incubated for 2.5 hrs, maintained in prophase I, and then imaged for 9 hrs with a sampling frequency of 15 mins. The data are reported as YPet signal corrected by averaged mCherry signal. Each point is the mean ± SEM of individual oocyte traces from three independent experiments and the total number of oocytes analyzed is in parentheses. Insertion of CPEs to any position in the 3'UTR of *Oosp2* was not sufficient to repress translation. **c)** Translation rates of wild type *Oosp1*, wild type *Oosp2*, and mutant *Oosp2* reporters in GV-arrested oocytes. The translation rate for each oocyte was calculated by linear regression of the reporter data (Supplementary Fig. 6b) between 6 and 9 hrs. The number of oocytes analyzed and mean ± SEM are reported. Statistical significance was evaluated by Kruskal-Wallis test; ****: $p < 0.0001$; ns: not significant.

### Supplemental Figure 7

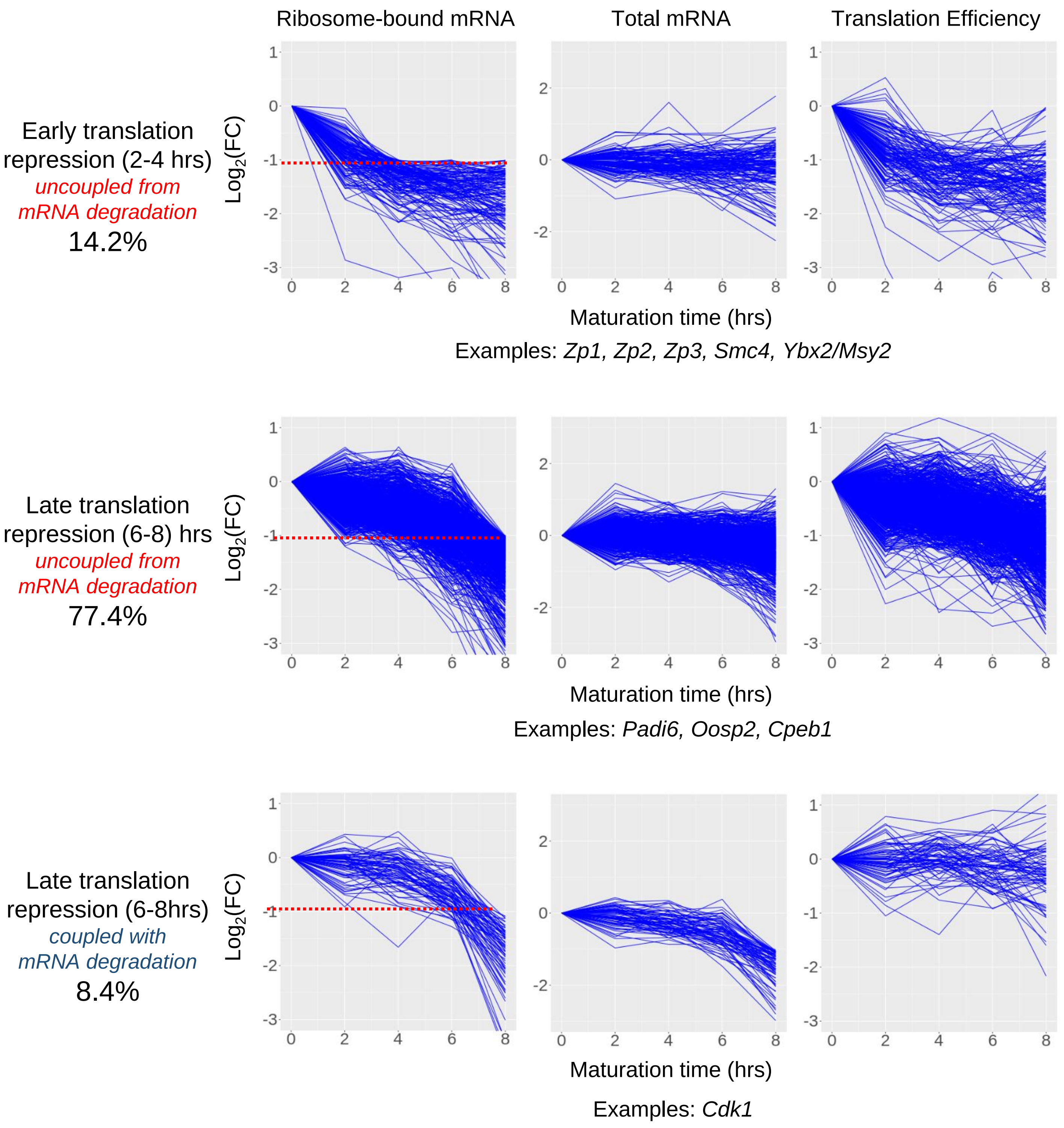

**Supp. Fig. 7. Progressive repression of translation during meiotic maturation**

Pair-wise statistical analysis of TMM-normalized CPM values was performed by comparing each point to 0 hrs. Messenger RNAs were classified as undergoing early translation repression if ribosome loading was significantly decreased between 2 to 4 hrs and remained decreased at 6 and 8 hrs (ribosome-bound mRNA: $LFC \leq 1$ and $FDR \leq 0.05$). Messenger RNAs were classified as undergoing late translation repression if ribosome loading was only significantly decreased between 6 to 8 hrs (ribosome-bound mRNA: $LFC \leq 1$, $FDR \leq 0.05$). The majority transcripts (86.1%) become translationally repressed without simultaneous changes in the total mRNA level (total mRNA: $1 \geq LFC \geq -1$). Only 8.4% of translationally repressed mRNAs show concurrent degradation (total mRNA, $LFC \leq -1$). Fold changes in TE are reported for each message.

### Supplemental Figure 8

## *Zp2*

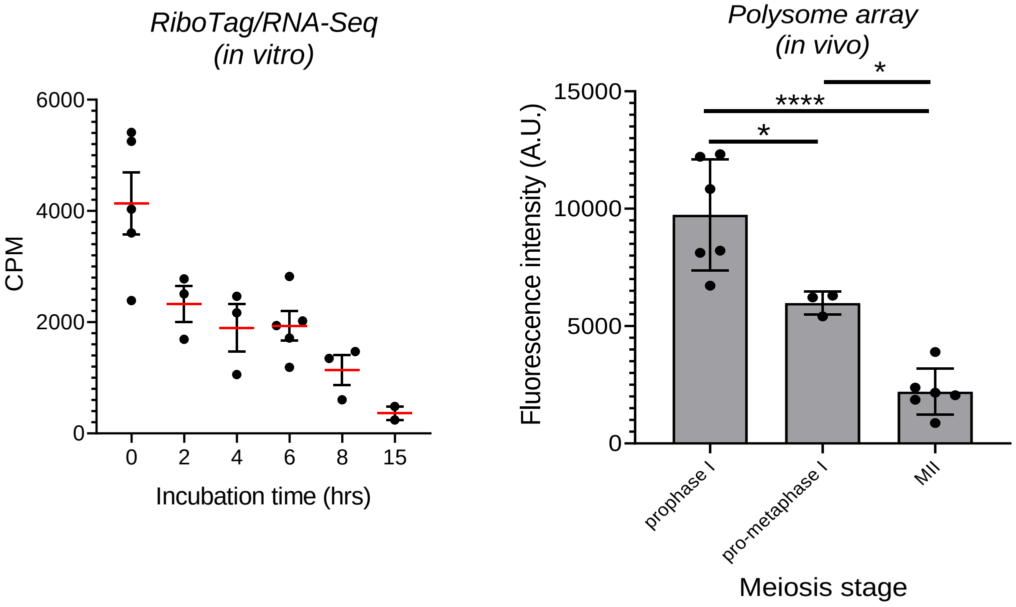


#### *Cpeb1*

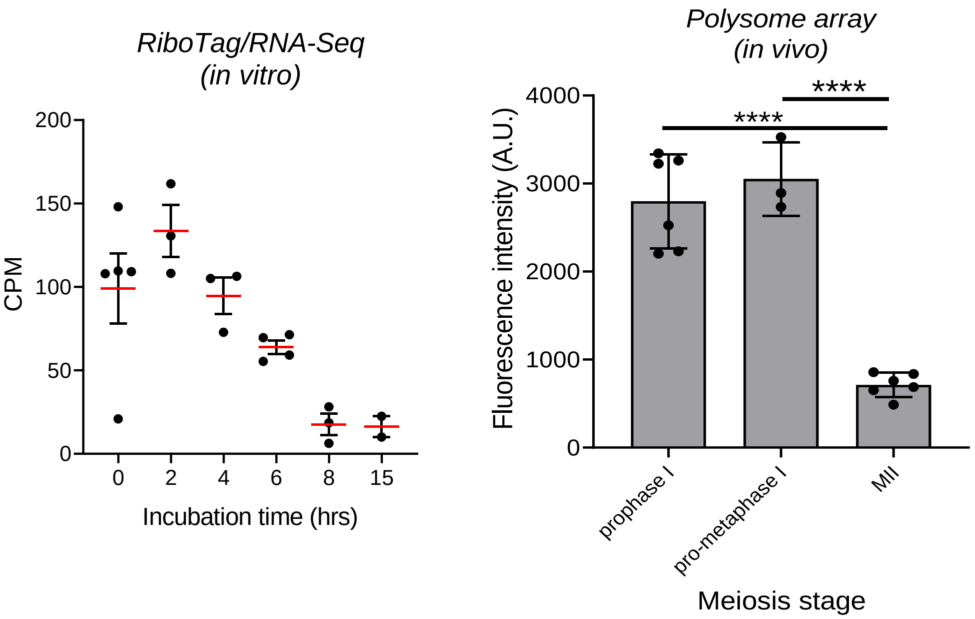

**Supp. Fig. 8. Translation repression is consistent across multiple biological replicates**

TMM-normalized CPM values from the two replicates of our RiboTag/RNA-Seq experiment were compared to a single point of a previous pilot RiboTag/RNA-Seq study[21], a 0 hrs/6 hrs dataset that is under review, a published GV/MI/MII dataset[42], and a published GV/MII dataset[27]. No cross study normalizations nor corrections for batch effects were applied. The translation of the *Zp2* and *Cpeb1* decrease progressively during meiotic resumption in all datasets both *in vitro* and *in vivo*. Note the difference in time course of translation repression for the two mRNAs.

### Supplemental Figure 9

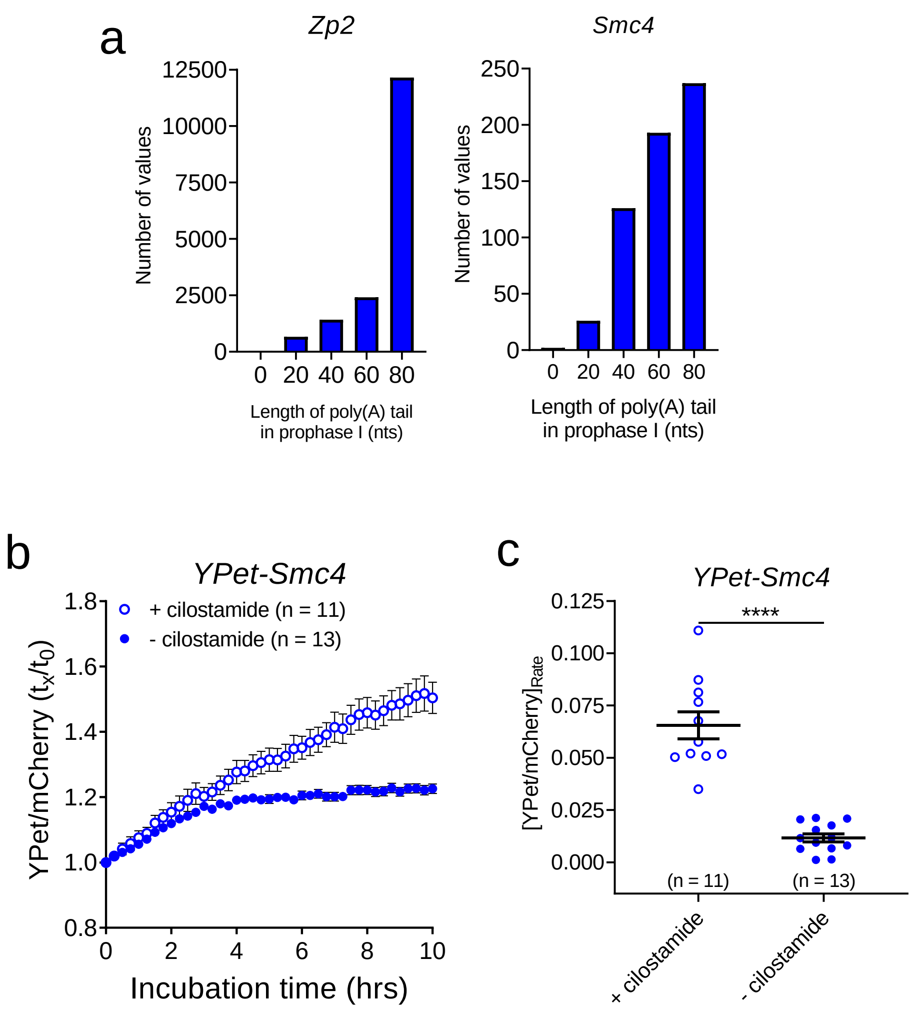

**Supp. Fig. 9. The 3'UTRs of *Zp2* and *Smc4* signal for rapid translation repression of the messages during meiotic maturation**

**a)** Polyadenylation state of *Zp2* and *Smc4* in GV-arrested oocytes. Mining of deposited TAIL-Seq data[29] reveals that 73% of *Zp2* transcripts have poly(A) tails with ≥ 80 nts and 40% of *Smc4* messages have poly(A) tails with ≥ 80 nts. These data are in agreement with the relatively high translation rates of these two mRNAs during prophase I arrest in our RiboTag/RNA-Seq dataset. **b)** Accumulation of *Smc4* reporter in GV-arrested or maturing oocytes. GV-arrested oocytes were collected, microinjected with *YPet-Smc4* along with polyadenylated *mCherry*, and allowed to recover for 16 hrs. Oocytes were then either maintained in prophase I arrest with cilostamide treatment (empty circles) or allowed to mature (solid circles) and imaged for 10 hrs with a sampling frequency of 30 mins. Data are reported as the fold change of the YPet/mCherry ratios as compared to 0 hrs. Each point is the mean ± SEM of individual oocyte traces obtained in a single experiment. The total number of oocytes analyzed is in parentheses. **c)** Translation rates the *Smc4* reporter in GV-arrested or maturing oocytes. The translation rate for each oocyte was calculated by linear regression of the reporter data (Supplementary Fig. 9b) between 6 and 9 hrs. Mean ± SEM is reported. Statistical significance was evaluated by Mann-Whitney test; ****: $p < 0.0001$.

### Supplemental Figure 10

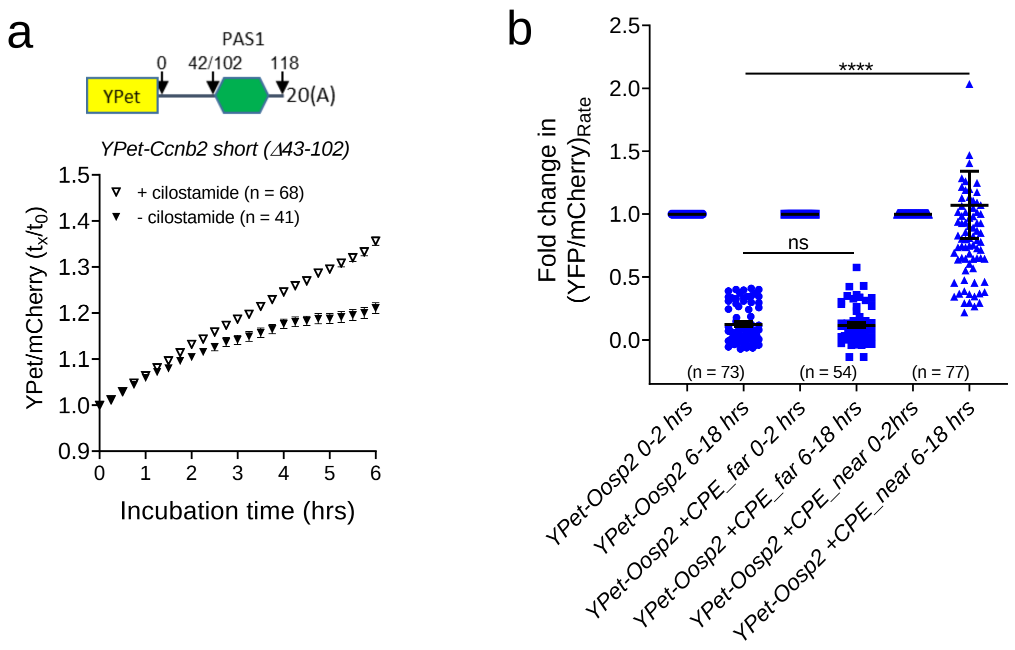

**Supp. Fig. 10. Translation repression occurring during meiotic maturation requires a CPE near the PAS in the 3'UTR**

**a)** Accumulation of a mutant *Ccnb2* reporter in GV-arrested or maturing oocytes. A mutant 3'UTR (*YPet-Ccnb2 short (Δ43-102)*) was obtained by deleting the CPE as well as the sequence downstream of the binding element but upstream of the PAS. GV-arrested oocytes were collected, microinjected with *YPet-Ccnb2 short (Δ43-102)* along with polyadenylated *mCherry*, and allowed to recover for 16 hrs. Oocytes were then either maintained in prophase I arrest with cilostamide treatment (empty circles) or allowed to mature (solid circles) and imaged for 10 hrs with a sampling frequency of 15 mins. Data are reported as the fold change of the YPet/mCherry ratios as compared to 0 hrs. Each point is the mean ± SEM of individual oocyte traces obtained in two independent experiments. The total number of oocytes analyzed is in parentheses. **b)** Translation rates of wild type *Oosp2* and mutant *Oosp2* mutants reporters in maturing oocytes. The translation rate for each oocyte was calculated by linear regression of the reporter data (Fig. 7f) between 0 and 2 hrs or 6 and 18 hrs. Mean ± SEM is reported. Statistical significance was evaluated by Kruskal-Wallis test; ****: $p < 0.0001$.

### Supplemental Figure 11

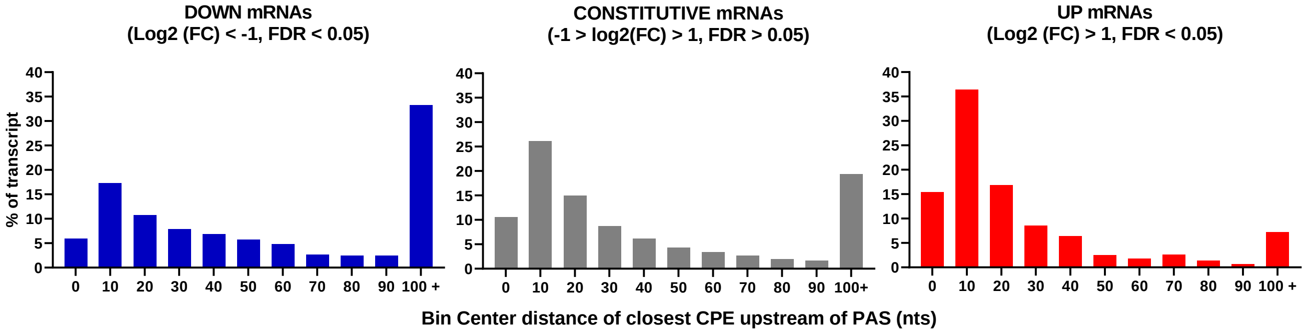

**Supp. Fig. 11. The presence of a CPE proximal to the PAS prevents translational repression during meiotic maturation**

Using deposited 3'UTR sequences, the distance between each PAS and each CPE was calculated for every message present in our RiboTag/RNA-Seq dataset. The distance of the closest CPE to a PAS was then calculated and plotted for UP, DOWN, and CONSTITUTIVE mRNAs. This analysis shows that messages whose translation did not decrease during meiosis (grey and red) have an enrichment of CPEs within 30 nts upstream of the PAS.

### Supplemental Figure 12

Early translation activation (2-4 hrs)
30.0%

Late translation activation (6-8 hrs)
70.0%

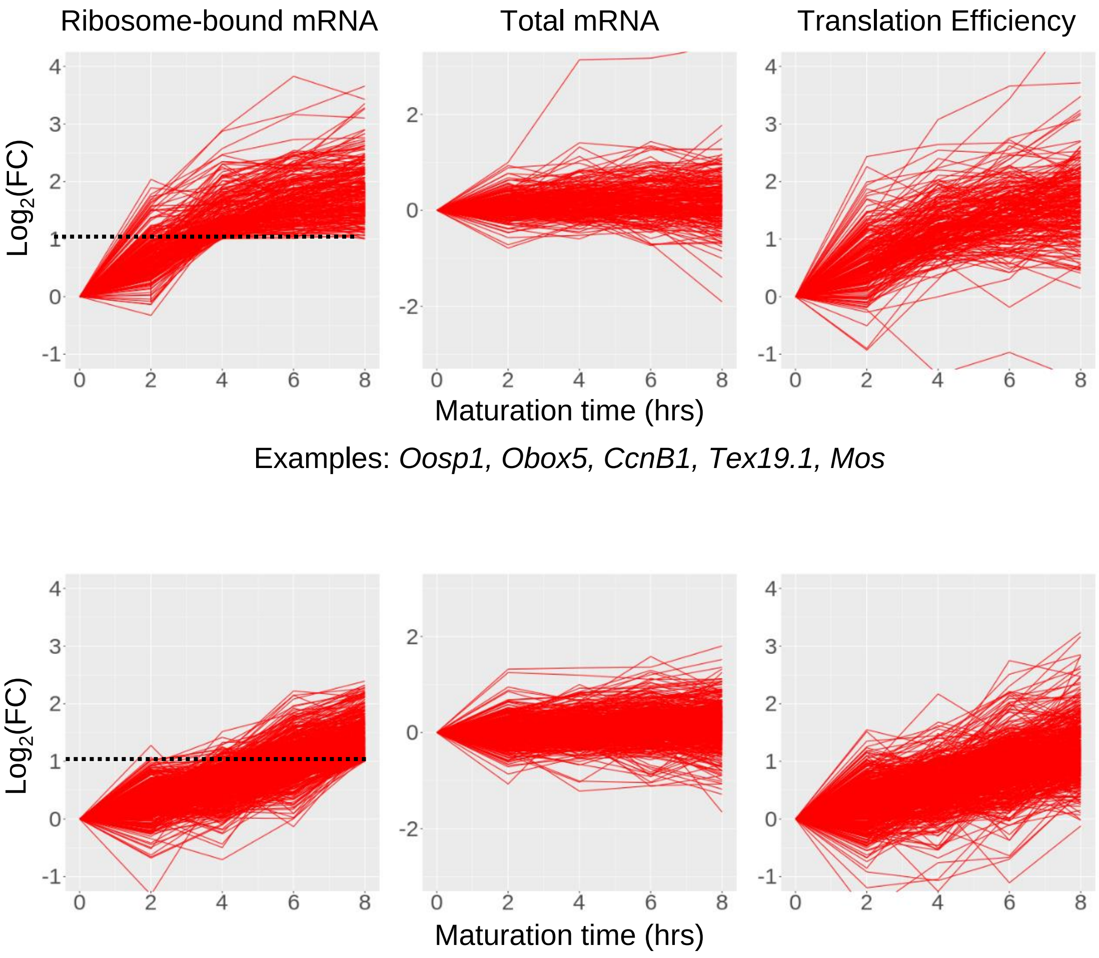


Examples: *Oosp1, Obox5, CcnB1, Tex19.1, Mos*

Examples: *Oosp3, Dazl, Cnot9, Cpeb4, Btg4*

**Supp. Fig. 12. Progressive activation of translation during meiotic maturation**

Pair-wise statistical analysis of TMM-normalized CPM values was performed by comparing each point to 0 hrs. Messages were classified as undergoing early translation activation if ribosome loading was significantly increased between 2 to 4 hrs and remained increased at 6 and 8 hrs (ribosome-bound mRNA: LFC ≥ 1 and FDR ≤ 0.05). Messages were classified as undergoing late translation activation if ribosome loading was only significantly increased between 6 to 8 hrs (ribosome-bound mRNA: LFC ≥ 1 and FDR ≤ 0.05). Fold changes in TE are reported for each message.

### Supplemental Figure 13

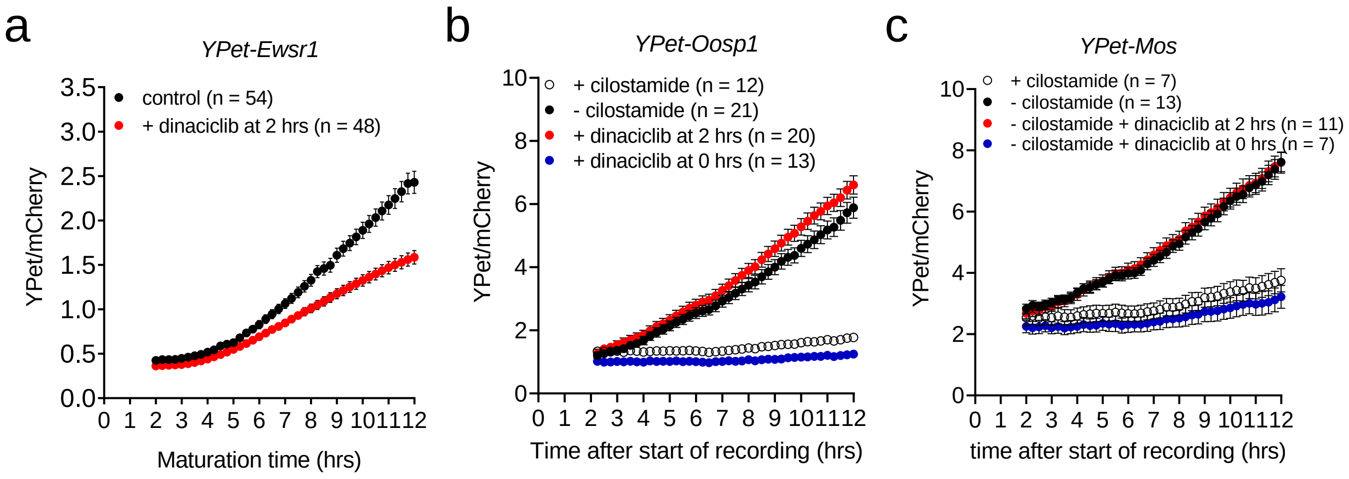

**Supp. Fig. 13. CDK1 inhibition differentially affects the translation activation of activated maternal messages**

The effect of CDK1 inhibition on the translation of activated maternal messages, **a)** *Ewsr1*, **b)** *Oosp1*, and **c)** *Mos*. GV-arrested oocytes were collected and microinjected with oligo-adenylated *YPet-Ewsr1*, *YPet-Oosp1*, or *YPet-Mos* mRNA along with poly-adenylated *mCherry* mRNA. Two groups of oocytes were maintained in prophase I arrest with cilostamide treatment (empty, black circle) or were incubated with a CDK1 inhibitor (dinaciclib) without cilostamide at the start of maturation (blue circle). Another two groups of oocytes were matured without (black circle) or with dinaciclib added at 2 hrs after release (red circle). Imaging started 2 hrs after cilostamide release and lasted for 10 hrs with a sampling frequency of 15 mins. Each point is the mean ± SEM of individual oocyte traces obtained in three separate experiments. The total number of oocytes analyzed is in parentheses.

### Supplemental Table 1

| Gene | 3'UTR sequence |
|---|---|
| *Ccnb1* | ctccaatagactgctacatctgcagatgcagttggcaccatgtgccgcctgtacataggatacctaccgtgtttacttgctcttcaataaaggtt<br>gtgacttctcattttacatagcttaactcatttgaatgttgttgcttctgagtttaggctaacggaagttgtcgaatttaggagtatattaaaaactgc<br>atctagttttaacagtggatccaactaatgtatatatctgtagcctatatgtctatatacatccttcactgtgtgtccttatatcatcatgtcttctgcct<br>cactctagtttaaactctaaatctaccagctagtcctttgttccattttccagtggttgccacctttaaccactgtctcttggtttgtcaactttcagat<br>ctgaaaccaagtatctttttttatgtaattatttatttgttcttaattggaaaataggatgttcaaaattaaaggtgtgttttaaaaagaatttgccccc<br>aagtctcactatcaacagataagggtgtattcttgtatatcctgtatagatataatcatgcatatactcccaaggagatatttttatatgggttcatt<br>ttatcaacagtattcctatcagcattcctttcaatgcctatattgcatttcctagtgtgaacaaactgtgtgtaacatagtcattccctcggtgggat<br>tcaagtgcattctctcagtgccctccacagtgttcttaaatgatgtttaatgtcttgcttggcttcattcatagtagctcttccaggggtgtgctttga<br>attctgacagccagatgggtgtggctgccaccataccaaggcgccactcctgtcttgtaatgccacctggaaaagaatcctgtctcatttgct<br>gttttaatttatacatctgatatcaagttgaataaaatttattggtggaaagctt |
| *Ccnb2 short* | gcactggactgccatcctgcgcttctcagatcctgtatgtattttattctagtttacatcacaaacctcttctcagactcattttctaattgtgtattgat<br>gaaaaataaagctattg |
| *Ewsr1* | agacctgcagagctgcattgacgaccagatttatttttaaaccagaaaatgttttaaatttataattccatatttataatgttggcgacaacatta<br>tgattattccttgtctgtactttagtattttcaccatttgtgaagaaacattaaaacaagttaaatagtagtgtgctgagttttttttgttgttgtttttt<br>gatttgttcttttttttttcctttttaagatggttgttttaagacttaacaatgggaaccccttgtgagcatgctcagtatcattgtggagaaccaagag<br>ggcctctaattgtaacaatgttcatggttgtgatttttttttttaaataaaattccaaatgttt |
| *Mos* | ctccatcgagccgatgtagagataagctttttgtctctgtttatttttttaaagaagtaaggatggtgtggaagaaaacataccactagggcata<br>tttttaggaaataaagttaccacgaacttc |
| *Oosp1* | atggtctggtgatttcttatctcctttttaaaaagatacaattttcaattttaaataaagtaaaatgcgaccttcatatattctcatttttaaaaaggag<br>ctgtccctgcttttaattgtataaaatctgattaaagaataaaaatttcccattagtgccgtgtctaa |
| *Oosp2* | ggacagtgctggcaccagagatttcactgtaagacaaacttttctctttcaataagaaaaaaaatcctggcataagcttctccttagagatctt<br>attcctatacatcaatgaaataaaggctcagcttatgaagatgca |
| *Smc4* | agttgttcacactgaaaattttcaaatctagattgggtgttctagttttgtctgtaaaagactgtaagttgtataaaatacatttttctaaattg |
| *Zp2* | ttggacttgcaaataaagagactgcagtt |

#### Supp. Table 1

Sequences of the wild type 3’UTRs used to construct YPet reporters. These sequences were obtained from the BAM data of the RiboTag/RNA-Seq experiment.

#### Data availability

Raw sequences and TMM-normalized CPM values from the RiboTag/RNA-Seq experiment will be available on UCSF Box.

#### Code availability

Scripts used for statistical analysis of the RiboTag/RNA-Seq data and those used to calculate CPE and PAS distances will be available on UCSF Box.
